# Achieving reproducibility and accuracy in cancer mutation detection with whole-genome and whole-exome sequencing

**DOI:** 10.1101/626440

**Authors:** The Somatic Mutation Working Group of the SEQC-II Consortium

## Abstract

Clinical applications of precision oncology require accurate tests that can distinguish tumor-specific mutations from errors introduced at each step of next generation sequencing (NGS). For NGS to successfully improve patient lives, discriminating between true mutations and artifacts is crucial.

We systematically interrogated somatic mutations in paired tumor-normal cell lines to identify factors affecting detection reproducibility and accuracy. Different types of samples with varying input amount and tumor purity were processed using multiple library construction protocols. Whole-genome and whole-exome sequencing were carried out at six sequencing centers followed by processing with nine bioinformatics pipelines to evaluate their reproducibility. We identified artifacts due to sample and library processing and evaluated the capabilities and limitations of bioinformatics tools for artifact detection and removal.

By examining the interaction and effect of various wet lab and computational parameters concomitantly, here we recommend actionable best practices for mutation detection in clinical applications using NGS technologies.

## Introduction

The outlook for cancer patients in desperate need of new medicines, while improving, is still not encouraging - oncology drugs in clinical trials have only a 6.7% likelihood of approval^1^. Another crisis looms in the pre-clinical research space, where various studies have estimated irreproducibility rates of 51%-89%^2–4^. Based on these rates, approximately US$28 billion/year is spent on irreproducible pre-clinical research in the US alone^5^ which could be spent on furthering the development of better medicines to treat patients in need. While the imperative for more accurate translational research and precision medicine is clear, currently available techniques, methods, and tools are still confounded by reproducibility issues.

In precision oncology, identifying whether a patient carries specific genetic aberrations relevant to a treatment or disease can guide pharmaceutical target identification, inform patient therapy, dictate dosing, suggest probability of adverse events^6^, and/or determine membership of an individual into a specific patient subset. As the cost of NGS drops precipitously, more researchers and clinical lab members are turning to this tool to identify mutations in clinical samples. Currently, a dizzying panoply of sample processing protocols, library preparation methods, sequencing machines, sequencing centers, and bioinformatics pipelines exist. However, there are no comprehensive studies on how each step influences the outcome when used with various combinations of tools and technologies. Moreover, samples can arrive at the testing lab in different states (fresh vs FFPE), the amount of input DNA can be variable, and tumor purity is rarely consistent across clinical samples.

Previous studies have addressed individual components of somatic variant calling in isolation; for example bioinformatics pipelines^7,8^ or sample factors^9^ have been studied individually. To date, no study has addressed the effect of cross site reproducibility together with the potentially influential interactions between biological, technical, and computational factors on the accurate identification of variants. The overarching goal of this work is to develop systematic methods for evaluating performance using realistic reference datasets. Here we profile well characterized^10–12^ and readily available paired cell lines (breast cancer and matched normal) with different types of biosample, input amounts, library preparations, sequencers, sequencing centers, and bioinformatics pipelines to thoroughly investigate how these factors and interactions between them may affect mutation detection. Based on our findings, we present actionable best approaches throughout the sequencing and analysis processes to enhance reproducibility, accuracy, specificity, and sensitivity.

## Results

### Study design

To date, great efforts have been devoted toward seeking an optimal strategy for detecting cancer mutations using NGS^13–19^. Most studies have compared bioinformatics pipelines/callers based on the accuracy and consistency of mutation detection^17–19^, while only a few studies have compared other components such as assay development^15^, library preparation^14^, and biosample resources^20,21^. Most of these studies utilized *in-silico* approaches with simulated ground truth or real tumor samples^13,15,16,22^ for benchmarking. The data sets or samples from previous studies either do not accurately represent real human tumor biopsies or are not sustainable for multiple benchmark studies. To our knowledge, only one attempt has been made to characterize a cancer cell line (from metastatic melanoma) for the purposes of creating a long-term community reference sample. However, the scale of that study was limited and did not provide extensive analysis on the performance of different NGS platforms^23^.

To pinpoint factors affecting somatic variant calling, a matched pair of breast cancer cell lines (tumor HCC1395 and normal HCC1395BL) was selected for profiling. Previous studies of this triple negative breast cancer cell line have indicated the existence of many structural rearrangements and ploidy changes^24^. Several attempts have been made to identify these structural variations and copy number changes, in addition to single nucleotide variation (SNV) and small insertions/deletions (“indels”)^11,12,25^. Given that appropriate consent from the donor patient has been obtained for tumor HCC1395 and normal HCC1395BL for the purposes of genomic research, we sought to characterize the pair of cell lines as possible reference samples for the NGS community. To fully characterize all components and aspects of variant calling together in one study, we designed and conducted a series of experiments to capture as many variables as possible across the whole process of NGS cancer mutation detection including: biosample preservation differences, amount of input DNA, library preparation protocol, NGS sequencing platforms, tumor purity, read coverage, and bioinformatics analysis pipelines.

To compare the tumor and normal samples, bulk DNA was extracted from fresh cell cultures. In addition, to mimic FFPE sampling, cell pellets were suspended in a small volume with the addition of histogel. The cell “blocks” were treated with formaldehyde for different durations (1, 2, 6, and 24 hours) and then embedded in paraffin for storage. DNA from “FFPE” blocks was extracted based on a standard protocol (see Supplemental Methods). Lastly, to evaluate the effects of variable tumor purity, we combined tumor DNA with normal DNA at different ratios to create a range of mixtures representing tumor purity levels of 75%, 50%, 20%, 10% and 5% (**Suppl. Fig. 1**).

Illumina TruSeq PCR-free library preparation was used for all sample preps with input amounts over 100 ng, and lower inputs (<=100 ng) were prepared with either TruSeq-Nano or Nextera Flex. Whole-genome sequencing (WGS) for all samples and whole-exome sequencing (WES) for a subset of samples were performed across different sites. Sequencings were performed using different HiSeq platforms (HiSeq1500, HiSeq2500, HiSeq4000, and HiSeqX), as well NovaSeq, the latest Illumina high throughput sequencer. (**Fig. 1a**).

**Figure 1.**
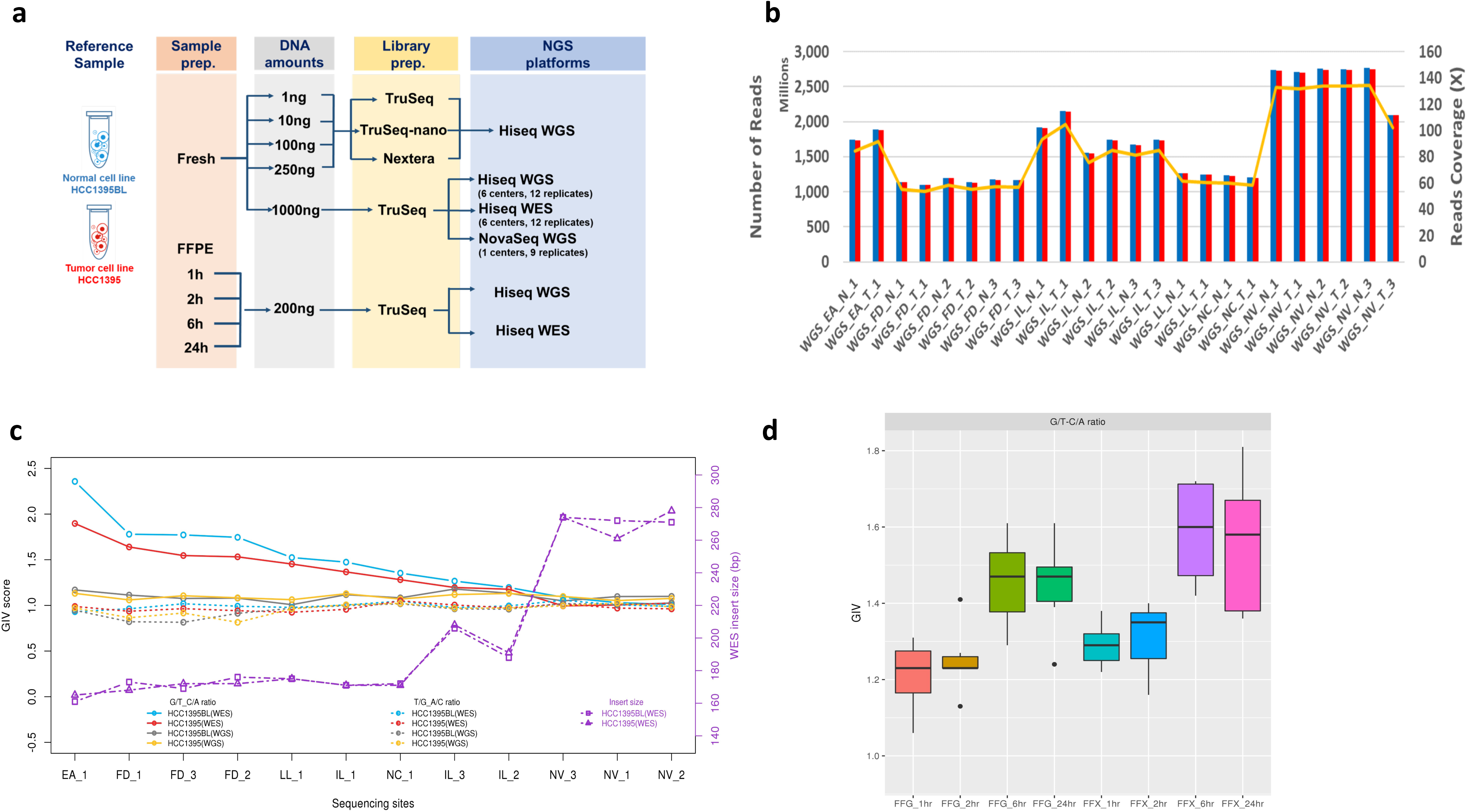
Study design and read quality. **(a)** Study design to capture “wet lab” factors affecting sequencing quality. DNA was extracted from either fresh cells or FFPE processed cells (formalin fixing time of 1, 2, 6, or 24 hours). Both fresh DNA and FFPE DNA were profiled on WGS and WES platforms. For fresh DNA, six centers performed WGS and WES in parallel following recommended protocols with limited deviation. Three of the six sequencing centers generated library preparation in triplicate. For FFPE samples, each fixing time point had six blocks that were sequenced at two different centers. Three library preparation protocols (TruSeq PCR-free, TruSeq-Nano, and Nextera Flex) were used with four different quantities of DNA inputs (1, 10, 100, and 250 ng). All libraries from these experiments were sequenced on the HiSeq series. In addition, nine libraries using the TruSeq PCR-free preparation were run on NovaSeq for WGS analysis. **(b)** Read yields (blue) and mapping statistics (red) from twelve repeated WGS runs. **(c)** Global Imbalance Value (GIV) of G>T/C>A and T>/A>C mutation pairs in WES and WGS runs. Six centers used a range of time (80 to 300 seconds) for DNA sharing. As a result, insert DNA fragment sizes ranged from 161 bp to 274 bp. **(d)** Distribution of GIV scores of FFPE DNA with four different fixing times (1, 2, 6, and 24 hours), analyzed with WES or WGS. FFX: FFPE on WES platform, FFG: FFPE on WGS platform.

### Survey of read quality

WGS was performed at six sequencing centers, generating a total of 42 sequencing results from the standard TruSeq PCR-free libraries prepared from 1000 ng input DNA. There were three different HiSeq platforms in the cross-platform comparison including HiSeq4000, HiSeq X10, and NovaSeq. All sequencing centers and platforms produced high quality data as demonstrated by base call Phred quality scores above Q30, and greater than 99.8% of reads mapped to the reference genome (GRCh38). Variation was observed in the quantity of reads per sample generated, with some centers consistently delivering much higher coverages (100X) than others (50X), which was driven by sequencing platform yield differences as well as run pooling schemes and coverage targets (**Fig. 1b**). Among the WGS libraries prepared using fresh cells, insert size distribution and G/C content was uniform (40 – 43% G/C). All of the WGS libraries had very low adapter contamination (<0.5%). Less than 10% of reads mapped redundantly for most libraries, indicating the high complexity of the WGS libraries. Similar mapping statistics were observed in NovaSeq WGS runs (**Suppl. Table 1**).

Similarly, WES performed across six sequencing centers using three different HiSeq models (HiSeq1500, HiSeq2500, and HiSeq4000), generated sequencing results where 99% of reads were mapped successfully (**Suppl. Table 2**). The largest variations in sequencing yield and coverage on target were seen between sequencing centers and even sometimes between different replicates of the same cell line at the same sequencing center. The variations were largely due to uneven library pooling, and sequencing yield differences between platforms. For the WES libraries, we observed much higher adapter contamination among all six centers, compared with WGS data. More than 20% of sequences in the WES from two sequencing centers came from the adapters while the rest of the centers had less than 12% adapter contamination (**Suppl. Fig. 2a)**. The higher adapter contamination was related to the longer sequencing read length in base pairs (bp) used (2× 150 bp vs. 2×125 bp). The WES libraries had higher G/C content (44 - 54%) compared to WGS libraries (40 – 43%). The libraries with more serious adapter contamination also had much higher G/C content compared to the rest of the WES libraries. Generally, regions with more reads on the target region had higher G/C content (**Suppl. Fig. 2b)**.

When comparing library preparation kits across different DNA inputs, both similarities and differences were noted. The average percentage of mapped reads ranged from 96% to 99.9% across TruSeq PCR-free, TruSeq-Nano and Nextera Flex libraries prepared with 250, 100, 10, or 1 ng of DNA input. However, the percentage of non-redundant reads was very low (<20%) for TruSeq-Nano with 1 ng input presumably due to PCR amplification of a low input amount of DNA (**Suppl. Fig. 2c**). Nevertheless, the percentage of non-redundant reads for 1 ng with Nextera Flex (also a PCR-based protocol) was reasonably good (∼70% and comparable to the performance of TruSeq-Nano 100 ng protocol). In addition, the overall G/C content was not affected by DNA input amount or library preparation kit. The Nextera Flex Library prep may be superior to TruSeq-Nano for lower input DNA amounts. The FFPE libraries were prepared from cells treated with formaldehyde at four time intervals and these samples yielded results with a high percentage of mapped reads and non-redundant read frequencies that were comparable to the results generated from WGS libraries prepared with fresh cells (**Suppl. Table 1).**

### Evaluation of DNA quality

The Global Imbalance Value (GIV) is a commonly used indicator of DNA damage^26^. We have used it here to monitor DNA quality in NGS runs. Indeed, we observed high GIV scores for the G>T/C>A mutation pair in WES runs of both tumor and normal cell lines from some centers. The GIV for G>T/C>A scores were inversely correlated with insert fragment size (**Suppl. Table 3)**, which is highly associated with DNA shearing time. Longer shearing time resulted in shorter DNA fragments. Insert fragment size and G/C content were also inversely correlated, suggesting increased off-target (non-exome region) reads when larger DNA fragments were sequenced. We observed high G>T/C>A scores when insert fragment sizes were between 160-180 bp; when insert fragment sizes were larger than 200 bp, we observed little or no imbalance (**Fig. 1c**). In contrast, we did not observe this imbalance in WGS runs **(Suppl. Fig. 2d)**, for which the insert sizes were normally larger than 300 bp (**Suppl. Table 1, Suppl. Fig. 3a)**. We did not observe such imbalance in other mutation pairs, such as T>G/A>C (**Fig. 1c**). Previous reports demonstrate that excessive acoustic shearing is the major modulator of 8-oxoG damage ^27^. Therefore, the high ratio of G>T/C>A observed in some WES runs was likely an artifact of oxidative DNA damage during the fragmentation process.

Formaldehyde also causes the deamination of guanine. Thus, the GIV of G>T/C>A would also be a good indicator of FFPE induced DNA damage^28^. Indeed, we did observe “dose” dependent GIV imbalance in WGS runs of FFPE samples. Samples with longer formaldehyde fixing time had higher GIV imbalance scores, which indicated that those samples likely sustained more severe DNA damage. WES of FFPE samples showed consistently higher GIV imbalance than WGS of FFPE samples, indicating that excessive shearing and formaldehyde fixing have an additive DNA damage effect (**Fig. 1d**). Taken together, these results indicate that WES is more sensitive to site-to-site library preparation variation than WGS. Our result therefore provided one line of evidence indicating that WGS may be more suitable for FFPE samples than WES.

### Reproducibility of cancer mutation detection

To assess the reproducibility of cancer mutation detection with WES and WGS, we performed a total of twelve repeats of the WES and WGS analysis of the tumor and normal cell lines carried out at six sequencing centers. We used three mutation callers (MuTect2^29^, Strelka2^30^, and SomaticSniper^31^) on alignments from three aligners (Bowtie2^32^, BWA^33^, and NovoAlign) to generate a total of 108 Variant Call Format (VCF) files from WES and WGS analyses separately. The Venn diagram is widely used to display concordance of mutation calling results from a small number of repeated analyses^18^; however, this type of diagram is not suitable for such large data sets. To address this challenge, we applied the “tornado” plot to visualize the consistency of mutation calls derived across aligners, callers, and repeated NGS runs. The number of SNVs unique to one single VCF file are represented by the width of the tornado at the bottom, and the number of SNVs called in all VCF files are represented by the width at the top. Thus, like the actual meteorological event, a tornado that is touching down is “bad” (many called variants are likely false positives), and a tornado with the majority of the data at the top is “better” (many common variants called across all conditions).

In this study, as shown at the top of each plot in **Fig. 2a**, all aligners demonstrate a substantial pool of calls which are agreed upon under every repeated WGS or WES run called by three callers (MuTect2, Strelka2, and SomaticSniper). Each aligner also generated a pool of unique calls (variants identified in only one sample) represented at the bottom of the tornado plot. Reproducibility was better for WGS than WES, as we observed relatively more aligner-specific variants at the bottom of the WES tornado plots. We did not observe significant differences among results from the three aligners. However, calling results from WES tended to have more inconsistent SNV calls (bottom of tornado) than those from WGS, indicating that WES results may be less consistent than WGS results **(Fig. 2a)**.

**Figure 2.**
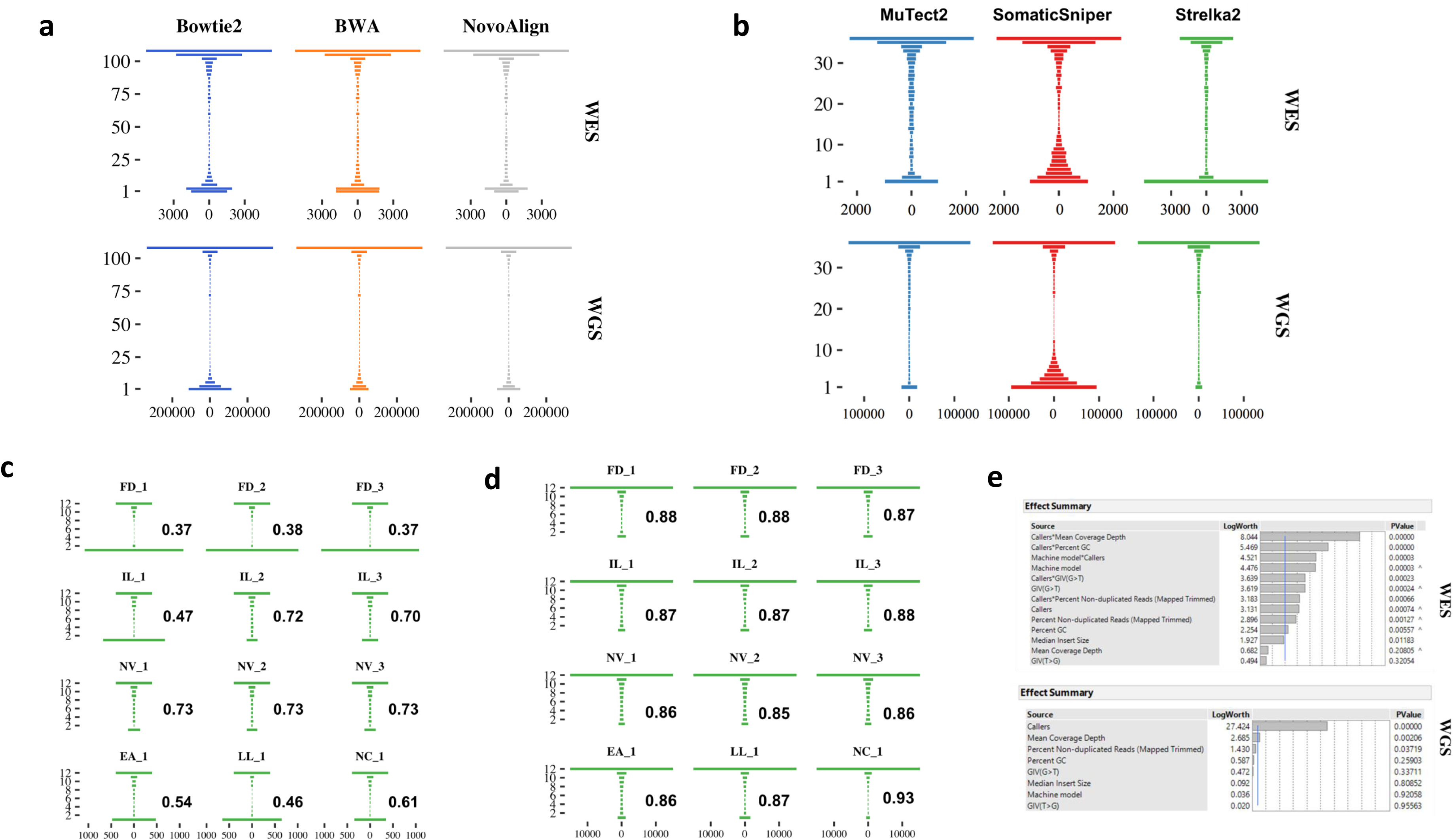
Mutation calling reproducibility. (**a**) “Tornado” plot of SNV overlaps across 108 VCF results from **twelve** repeated WGS and WES runs that were analyzed with **three** aligners, BWA, Bowtie2, and NovoAlign, and **three** callers, MuTect2, Strelka2, and SomaticSniper. The height of the “Tornado” represents the number of overlapping VCF files. The width of the “Tornado” represents the number of **accumulated** SNVs in that overlapping category. (**b**) “Tornado” plot of SNV overlaps across **36** VCF results from **twelve** repeated WGS and WES runs analyzed by MuTect2, Strelka2, and SomaticSniper from only the BWA alignments. **(c)** and **(d)** show the “Tornado” plots for **each of the 12** repeated WES and WGS runs, **respectively**, analyzed by Strelka2 from BWA alignments. The number in each plot represents the O_Score reproducibility measurement. (**e**) Effect summary of the model for WES or WGS. Effect tests were performed to evaluate the importance of each independent variable in a fixed effect linear model fitted for WES or WGS. F statistics and the corresponding P-value were calculated for the variables. Both P-value and LogWorth (-log10 (P value)) are plotted.

We then fixed the alignment to the community-accepted Burrows-Wheeler Aligner (BWA), and compared the results from our three callers, observing the differences in WES versus WGS run performance. While SomaticSniper displayed results that showed a very high similarity of tornado shape between WES and WGS analyses, those derived from the use of MuTect2 and Strelka2 showed greater divergence. The latter, especially Strelka2, were strikingly more consistent when used for WGS rather than WES (**Fig. 2b)**. Further examination of results from Strelka2 on BWA alignments of twelve repeated WES and WGS runs confirmed this observation (**Fig. 2c, d)**. Here we also introduced the O_Score, a metric to measure reproducibility of repeated analyses (see Supplemental Methods). O_Scores for Strelka2 and MuTect2 for WES runs were not only significantly lower than WGS runs, but also more variable. Unexpectedly, even though its overall O_Score was much lower than Strelka2 and MuTect2 in WGS runs, SomaticSniper showed better consistency in calling results in WES than in WGS (**Suppl. Fig. 4a, Suppl. Fig. 5)**. We also compared WGS runs with HiSeq vs WGS runs with NovaSeq using BWA for alignment and Strelka2 for calling. Both platforms were remarkably similar in terms of reproducibility, indicating that results from HiSeq and NovaSeq are comparable (**Suppl. Fig. 4b**). Taken together, Strelka2 had the best reproducibility in WGS repeated runs but the worst in WES runs, whereas MuTect2 had the best reproducibility in WES repeated runs **(Suppl. Fig. 4a, Suppl. Fig. 6).**

### Factors influencing reproducibility of cancer mutation detection

After establishing the O_Score to measure the reproducibility of NGS platform mutation detection, we also used it to determine what variables contribute most to variation between different repeated WGS or WES analyses separately. We included not only callers but machine model, read coverage, non-duplicated reads, G/C content, insert size, and two GIV scores (G>T/C>A and T>G/A>C). The combination of these parameters represented explained the majority of variation in WGS and WES runs, as we observed very high proportion of variance in the O_Score could be predicted by these variables. For WGS, the combination of these eight individual variables accounted for >99% of O_Score variance (R^2^ > 0.99) (**Suppl. Fig. 7a**). On the other hand, individual variables and five interaction terms (callers*coverage, callers*percent GC, caller*machine model, callers*GIV(G>T/C>A), and callers*non-duplicated reads) were significant for O_Score variance in WES runs (**Suppl. Fig. 7b**).

Overall, the most influential factor for the performance of WGS is the caller, followed by read coverage. As all twelve WGS runs were done with TruSeq PCR-free libraries with the same amount of DNA input, the percentage of non-duplicated reads did not affect reproducibility. While individual variables may influence WES run reproducibility, their effect levels were truly dependent on caller selection (**Fig. 2e**). Taken together, only a few factors (read coverage and callers) affect WGS reproducibility, whereas many factors, such as caller, reads coverage, insert fragment size, GC content, GIV (G>T/C>A) score, and their interactions influence WES run reproducibility. It is noteworthy that since an older machine model (HiSeq1500) was used only in one WES sequencing center, which also had low insert fragment size and high GIV (G>T/C>A) scores in sequencing reads, the impact of machine model on the WES run reproducibility could be a confounding factor.

### Definition of truth set for comparing and benchmarking

After comparing results within this study, we wanted to establish a truth set of mutation calls for the two cell lines. The schematic of mutation calling is displayed in **Fig. 3a**. In brief, we used BAMSurgeon^34^ to spike synthetic mutations into an HCC1395BL Binary Sequence Alignment/Map (BAM) file, matched with another HCC1395BL BAM file from the same sequencing center to create a synthetic tumor-normal pair. We then used the SomaticSeq^35^ machine learning framework with MuTect2^29^, Strelka2^30^, VarDict^36^, SomaticSniper, MuSE, and TNscope^37^ on the synthetic tumor-normal pairs to create sequencing- and aligner-specific classifiers that took into account error models within every sequencing center and aligner (**Fig. 3a**). These classifiers were used to generate high-confidence mutation call sets for tumor-normal pair of BAM files.

**Figure 3.**
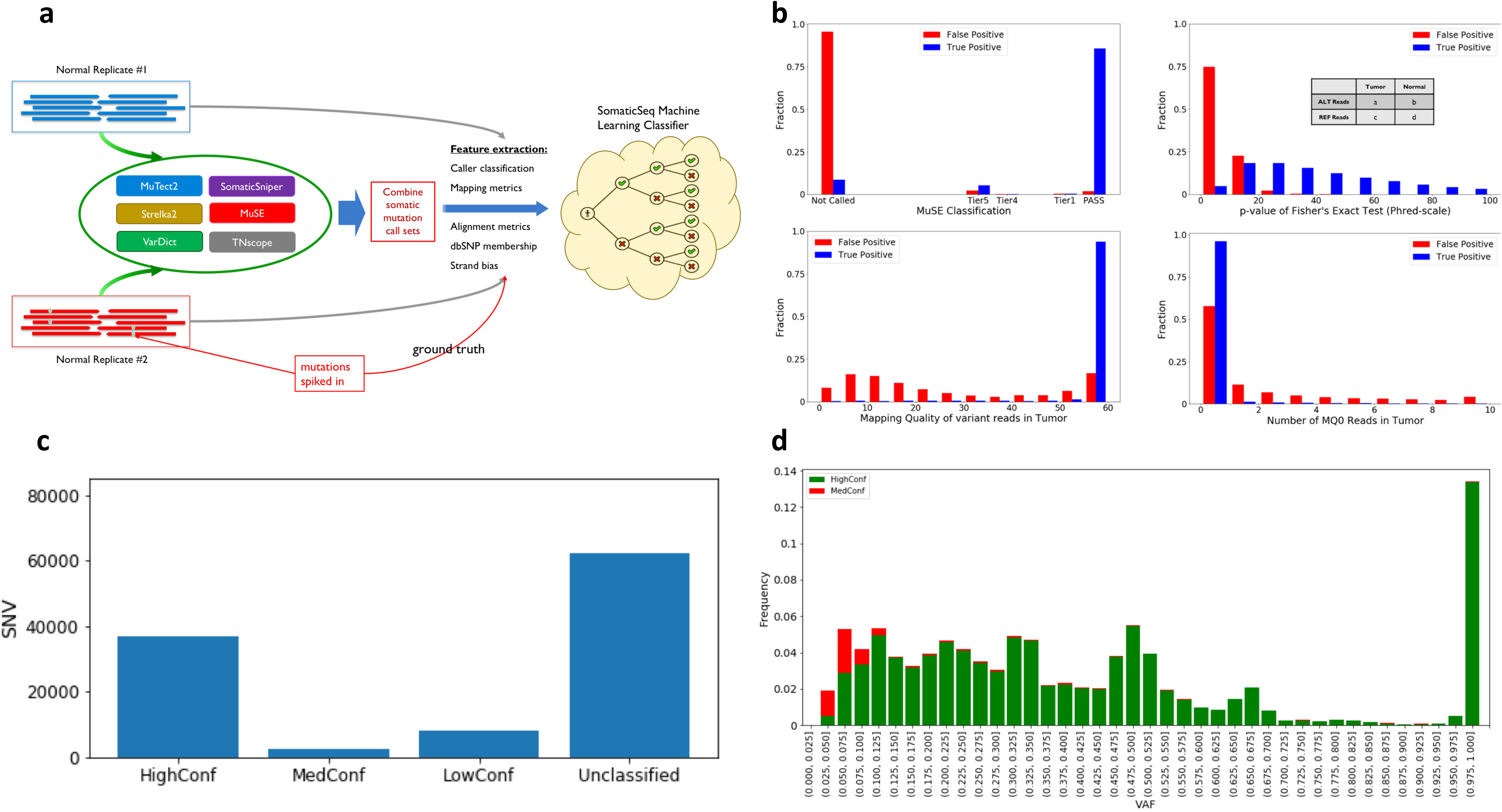
Definition of the truth set of somatic mutation calls. **(a)** General workflow for SNVs and indel truth set calling with 63 pairs of BAM files (21 repeated WGS runs on three aligners and six callers as specified in Fig. 1). **(b)** Four examples of 100+ features used in a building model for mutation calling. **(c)** Distribution of supporting evidence for called SNVs and indels. **(d)** VAF distribution of HighConf and MedConf SNVs in the truth set.

Next, factors driving either true positives or true negatives were investigated. The mapping quality of variant reads in the tumor was an important driving factor. For instance, for the vast majority of true somatic variant calls, the average BWA mapping quality (MQ) score for variant-supporting reads was around 60 (**Fig. 3b**). Variant calls with mapping qualities of 55 or below were overwhelmingly false positives. Similarly, a high number of MQ=0 reads also indicated false positives (**Fig. 3b**).

Finally, calls were categorized into four confidence-levels based on cross-aligner and cross-sequencing-center reproducibility, consistency with tumor purity titration data, and various red flags that are predictive of false positive status (**Fig. 3c**). The category with the highest number of evidence is labeled as “HighConf”. Those, along with variants labeled as “MedConf” are considered true positives in our truth set. The next category is labeled as “LowConf”, which has considerable, but often conflicting evidence. Thus, their status is ambiguous. The final category of “Unclassified” are calls that were deemed “PASS” in at least one sample, but were inconsistent across sequencing centers or aligners. These calls also had more samples that were classified as “REJECT” than “PASS”. Ultimately, we decided to use SNVs/indels in HighConf and MedConf categories as the truth set, which contained a total of 39,536 SNVs and 2,020 indels. In the truth set, there was a large fraction of seemingly homozygous calls (**Fig. 3d**). However, these homozygous calls might be misleading since karyotyping of HCC1395 indicated a large number of chromosomal aberrations. The ploidy of this cancer cell line is close to 3 with total of 66 chromosomes in truncated forms (**Suppl. Fig. 8a**). Therefore, we suspected that loss of heterozygosity (LOH) produced high levels of homozygous calls.

There were many SNVs/indels in our truth set with a variant allele frequency (VAF) less than 0.5. There were VAF peaks at 0.67, 0.5, 0.33, 0.25, and 0.125, which indicate different copy number states for those variants. Since most WGS runs defining the truth set were at less than 50X coverage (**Fig. 1b**), our lowest mutation allele frequency (MAF) variants were close to or slightly under 0.05. Thus, our truth set is incomplete. Nevertheless, based on the distribution of frequency versus MAF (**Fig. 3d**), we believe the number of true mutations missing in our truth set is very small and that our truth set should yield useful information to evaluate performance of tools and technologies.

### Effect of non-analytical factors on mutation calling

Before thoroughly investigating computational and bioinformatics analysis choices influencing mutation calling, we wanted to leverage our truth set to assess upstream non-analytical driver factors. First, we investigated the effect of read coverage by computationally varying the number of reads from the tumor sample and the normal sample. Across all three callers, true positives were increased by higher read coverage in both samples (**Fig. 4a**). Here, Strelka2 and SomaticSniper demonstrated sensitivity to low coverage of the normal sample, increasing the false positive call rate. Based on these data, 10X coverage of the normal sample is insufficient across all callers tested and >30X coverage for both tumor and normal samples is needed to lower the false positive rate. However, increasing coverage in the tumor sample did not translate to uncovering more true positive mutations. As a matter of fact, more false positives were observed when called by MuTect2 and SomaticSniper. This study was limited to 100% of tumor with 5% of MAF as the detection limit, which is an important caveat. To recover mutations with less than 5% MAF or in samples with less tumor purity, more coverage may be required.

**Figure 4.**
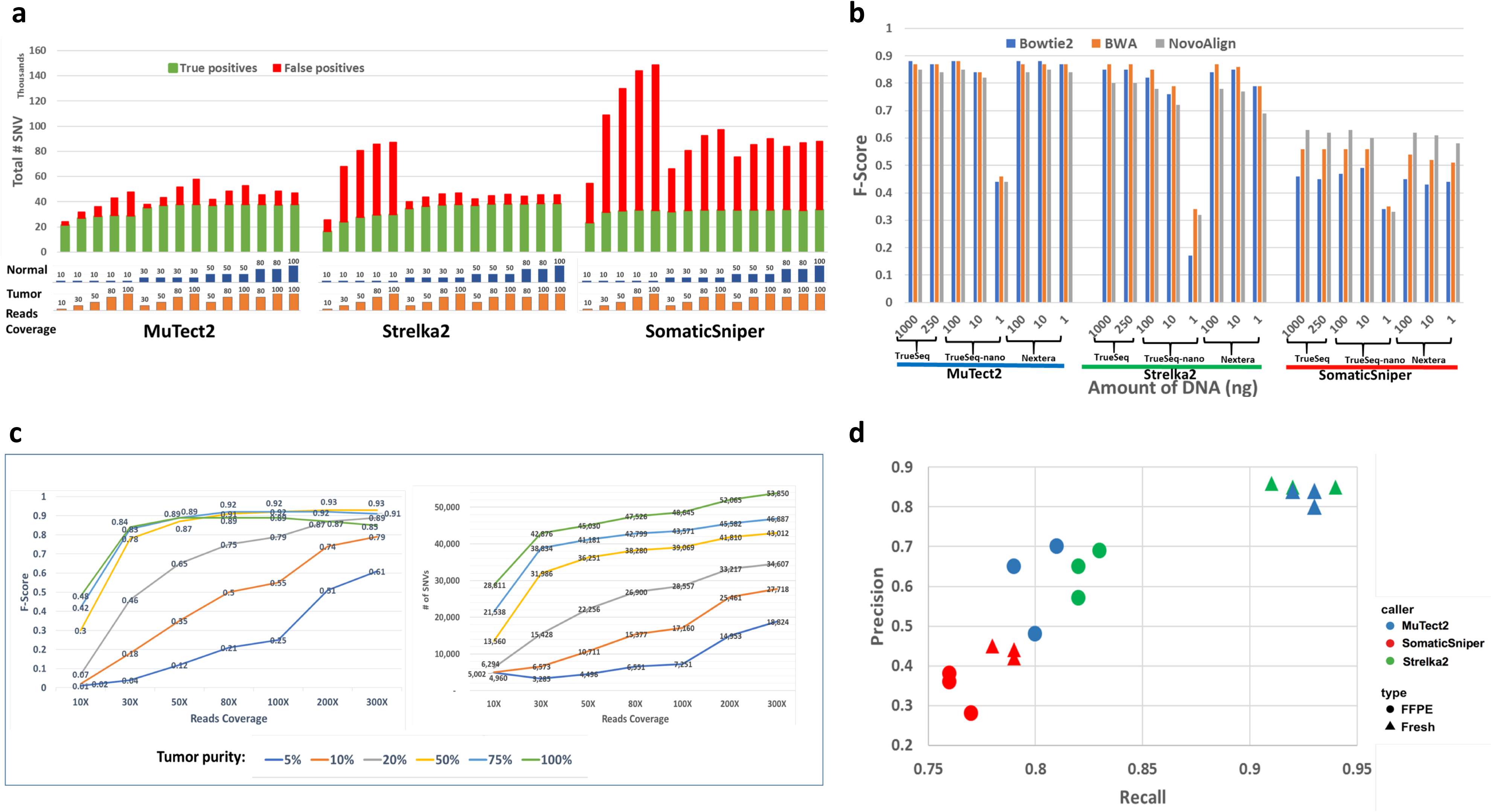
Non-analytical factors affecting mutation calling. **(a)** Effect of read coverage on mutation calling by three callers on BWA alignments. A pair of BAM files (tumor/normal) from BWA alignment with 100X read coverage was downsampled to generate a series of sub-files. MuTect2, Strelka2, and SomaticSniper were used to call mutations with various coverage combinations of the tumor/normal pair. SNVs from the truth set were used to define false positives and true positives by each caller at each coverage. **(b)** Performance of the callers on three library preparation protocols with different DNA inputs: 1, 10, 100, 250, and 1000 ng. **(c)** Performance of Strelka2 on BWA alignments with WGS runs on samples with various read coverages of samples with different purity levels. **(d)** Performance of MuTect2, Strelka2, and SomaticSniper on fresh DNA and FFPE DNA.

With three different library preparation protocols and varying DNA input amounts across multiple library preparations, we evaluated results from combinations of the three callers and three aligners. MuTect2 was found to be highly reliable, except for calling of the 1 ng TruSeq-Nano libraries (**Fig. 4b**). In contrast, Strelka2’s performance suffered severely from the 1 ng TruSeq-Nano libraries, as did SomaticSniper. These results demonstrate that 1 ng of DNA input in a TruSeq-Nano library preparation cannot be compensated by any aligner, caller, or aligner/caller combination tried here. Nextera Flex library preparation might be a better option for a low input DNA quantity.

Tumor purity is another non-analytical factor which can influence variant calling. Samples derived from human tumors contain not only tumor cells, but also a mix of non-malignant cells including immune cells, stromal cells, fibrotic cells, and others. The presence of these non-tumor cells likely skews variant calling results^17^. To explore this in a very simplistic approximation, a range of tumor purities was created by diluting tumor DNA into normal DNA at various ratios to mimic tumor purity at range from 5% to 100%, and coverages ranging from 10X to 300X were evaluated. Strikingly, for cases of very high tumor purity (75% or 100%), F-Scores actually dropped at the highest coverages, demonstrating the incompleteness of our truth set (>5% MAF) (**Fig. 4c**). Boosting read coverage may also have allowed more noise in and lowered performance since, as we observed, the total number of SNVs called continuously increased with more coverage. This is true not only for Strelka2, but also for the other callers (**Suppl. Table 4**).

Lastly, FFPE processing is another non-analytical factor known to have a major effect on variant calling results^38^. In this study, the same samples of fresh cells and cells processed with FFPE were called with Strelka2, SomaticSniper, and MuTect2. Both MuTect2 and Strelka2’s precision and recall were greatly reduced when samples were subjected to FFPE processing (**Fig. 4d**). On the other hand, SomaticSniper demonstrated only a small decrease in both metrics but otherwise underperformed significantly compared with the other two callers.

### Enhancing calling accuracy through bioinformatics

It is not always ideal to analyze challenging samples (FFPE or low tumor purity). However, that may be the only biosample available. In testing various informatics pipelines, some clearly overcome sample related biases more successfully than do others. To begin testing, samples were pre-processed using Trimmomatic^39^ or BFC^40^ to assess whether read trimming and error correction could affect recall or precision (as defined in Supplemental Methods) for variant calling of WES runs. Applying error correction with BFC to FFPE samples increased precision across most samples, but did not improve the recall rate (**Suppl. Fig. 9a**). However, precision is still improved for fresh DNA samples that have undergone BFC processing, but with a lower recall rate than results from Trimmomatic processing. Taken together, these results indicate that BFC is appropriate in cases of severe DNA damage, but may not be worthwhile if there is only mild damage from a process such as sonication.

FFPE is known to cause G>T/C>A artifacts as well^38^. Trimmomatic and BFC were investigated for their ability to detect these errors. Since the DNA damage causing the G>T/C>A mutation may not be isolated to only low quality base calls at the end of reads, Trimmomatic is not designed to remove this type of artifact. Trimmomatic processed data was more skewed toward C>A artifacts than was BFC processed data, which shows changes more broadly across nucleotide transitions (**Fig. 5a**). BFC reduced C>A artifacts, but introduced a small quantity of artifacts of other types, such as T>C mutations, indicating that caution should be exercised when using bioinformatics tools to correct FFPE artifacts.

**Figure 5.**
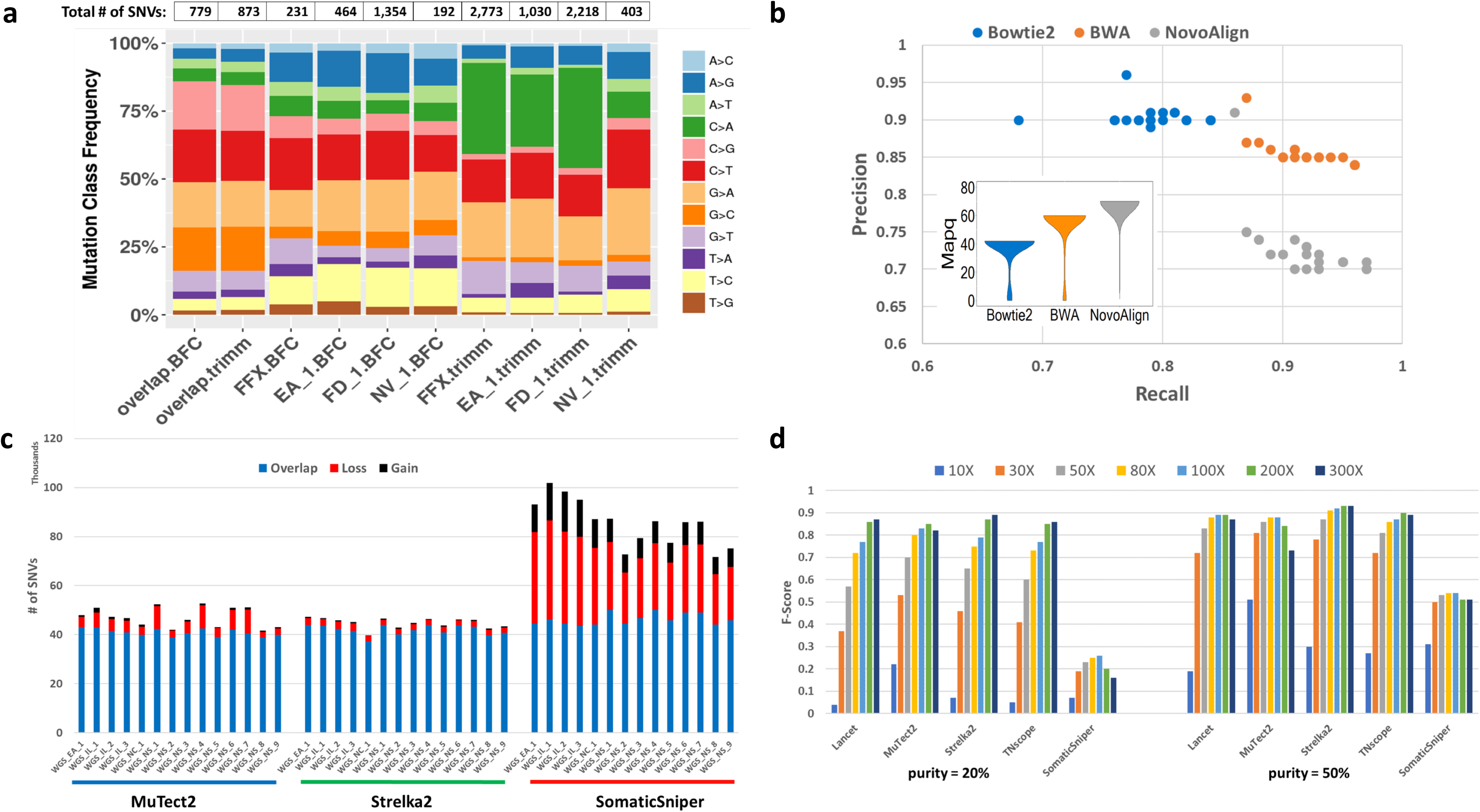
Bioinformatics for enhanced calling. **(a)** Distribution of mutation types called with Strelka2 on BWA alignment of four WES runs preprocessed by Trimmomatic or BFC. FFX: WES run on FFPE DNA, EA_1, FD_1, and NV_1, represented three WES runs with fresh DNA. **(b)** Performance of mutation calling by Strelka2 on three alignments, Bowtie2, BWA, and NovoAlign. The insert figure is a violin plot of mapping quality scores from three alignments for an example WGS run. **(c)** Effect of post-alignment processing (indel-realignment + Base Quality Score Recalibration [BQSR]) on mutation calling by MuTect2, Strelka2, and SomaticSniper. **(d)** Effect of tumor purity (20% vs 50%) on five callers: Lancet, MuTect2, Strelka2, TNscope, and SomaticSniper with read coverage from 10X to 300X.

Calling accuracy is dependent on the choice of caller and aligner, as well as how the caller and aligner interact. Strelka2 results with BWA-aligned reads were balanced, whereas results with Bowtie2-aligned reads seemed conservative. In contrast, Strelka2 results with NovoAlign-aligned reads seemed aggressive (**Fig. 5b**). When we examined the mapping quality scores for the three alignments, BWA’s mapping quality scores were usually between 50 and 60, Bowtie2’s scores were between 40 and 50, and NovoAlign’s scores were between 60 and 70. Strelka2 was trained and modeled on BWA alignment and thus works best in the bioinformatics context where it was developed^30^. Taken together these results indicate that there might be a joint effect between aligner and caller, depending on how callers were developed.

Next, the effect of the Genome Analysis Toolkit (GATK) post alignment and quality score recalibration was queried. MuTect2 and Strelka2 identified a very similar number of SNVs regardless of whether the process was employed (**Fig. 5c**). Conversely, SomaticSniper is highly sensitive to post alignment processing, with some SNVs gained and far more lost. Relatedly, the precision and recall rate change dramatically when this process was applied for SomaticSniper calling, but not for calling by MuTect2 or Strelka2 (**Suppl. Fig. 9b, c**). Clearly, our data confirm, with real-world experimental data, the importance of a full understanding of how the various components of mutation analysis by NGS methods work and interact with each other.

Tumor purity and coverage also play a role in caller performance. As expected, higher coverage can produce more SNV calls than does lower coverage (**Fig. 5d**). When tumor purity is high (>50%), 50X performs very similarly to 100X coverage across all callers tested. When tumor purity is low (<50%), calling is much more sensitive to sequencing depth. To test the performance of callers on low tumor purity samples, we pooled reads from the triplicate runs of WGS on samples that were sequenced at 100X coverage to generate coverage of 200X or 300X on each cell line. Besides the three main callers used in this study we also included two new tools (TNscope^37^ and Lancet^41^) to compare their capabilities in detecting mutations from a mix with as low as 5% of tumor DNA. With high tumor purity (>50%), we again observed that the accuracy of all the callers, with the exception of SomaticSniper, declined slightly with higher read coverage, indicating that our truth set lacked <5% MAF mutations. However, when tumor purity was 20% or lower, we observed that the benefit of higher coverage for mutation detection was realized. For a sample with 20% tumor, Lancet, Strelka2, and TNscope performed similarly well with 300X coverage (**Fig. 5d)**. On the other hand, SomaticSniper performed poorly at any tumor purity level, and increased read coverage did not rescue its performance. These results indicate that tumor purity is much more influential than coverage in the ranges tested here.

### Performance of WGS and WES across multiple sequencing centers

As this study leveraged six sequencing centers performing twelve WES and WGS experiments simultaneously, we were able to directly compare results from the two sequencing platforms within and across centers. To further understand if artifacts in WES library preparation are sequencing center-specific, we performed analyzed reproducibility of inter- and intra-centers based on concordance of SNV detection between any pair of NGS runs, in association of SNVs defined in the reference call set established in this study. Comparisons were carried out in four groups: a) WES (12 repeats/3 centers); b) WGS (limited to exome regions, 12 repeats/3 centers); c) WGS (12 repeats/3 centers), and d) WES vs WGS (limited to exome regions, 24 repeats/3 centers). We did each of these four comparison groups with “all reads” (original coverage) and “downsampled reads” (fixed coverage of each NGS run, 50X for WGS and 150X for WES). In addition to investigate reproducibility represented by overall SNVs called between a given pair of NGS runs, we also broke down SNVs into three subgroups: 1) In-truth set: SNVs defined in “HighConf” and “MedConf” categories; 2) Not in-truth set: SNVs defined in “LowConf” and “Unclassified” categories; and 3) Not defined: SNVs not defined in the reference call set. We then used average value of Jaccard scores for the concordance of SNVs in above three subgroups, to determine inter- and intra-center reproducibility for WGS and WES platform.

As shown in **Table 1**, results from WES were consistently less reproducible than WGS. When measured by overall called SNVs, inter-center variation was larger than intra-center variation. On the other hand, results from WGS, either limited to exome regions or the whole genome, were more consistent than results from WES. In addition, the difference in variation between inter-center and intra-center for WGS was very small.

**Table 1:** Cross-center/cross-platform mutation detection reproducibility with Strelka2: *Overall SNVs: all SNVs in a pair of NGS runs; In-truth set: SNVs defined in “HighConf” and “MedConf” categories; Not in-truth set: SNVs defined in “LowConf” and “Unclassified” categories; Not defined: SNVs not defined in above four categories.*

| Concordance of SNVs<br>(measured by Jaccard<br>index) |  | WES |  | WGS |  |  |  | WES vs WGS (Exome) |  |
| --- | --- | --- | --- | --- | --- | --- | --- | --- | --- |
|  |  | all<br>reads | downsampled | in-exome |  | whole genome |  | all<br>reads | downsampled |
|  |  |  |  | all<br>reads | downsampled | all<br>reads | downsampled |  |  |
| Overall<br>SNVs | overall | 0.36 | 0.29 | 0.80 | 0.74 | 0.74 | 0.69 | 0.47 | 0.40 |
|  | intra-center | 0.44 | 0.32 | 0.82 | 0.75 | 0.76 | 0.69 | 0.50 | 0.41 |
|  | inter-center | 0.35 | 0.28 | 0.79 | 0.74 | 0.74 | 0.69 | 0.47 | 0.40 |
| In-truth<br>set | overall | 0.90 | 0.83 | 0.92 | 0.88 | 0.91 | 0.88 | 0.89 | 0.82 |
|  | intra-center | 0.94 | 0.86 | 0.93 | 0.88 | 0.93 | 0.88 | 0.91 | 0.82 |
|  | inter-center | 0.89 | 0.82 | 0.92 | 0.88 | 0.91 | 0.88 | 0.89 | 0.82 |
| Not in-truth set | overall | 0.33 | 0.26 | 0.18 | 0.14 | 0.21 | 0.20 | 0.18 | 0.14 |
|  | intra-center | 0.44 | 0.30 | 0.22 | 0.15 | 0.25 | 0.21 | 0.21 | 0.15 |
|  | inter-center | 0.31 | 0.26 | 0.18 | 0.14 | 0.21 | 0.19 | 0.18 | 0.14 |
| Not defined | overall | 0.03 | 0.01 | 0.10 | 0.04 | 0.07 | 0.03 | 0.02 | 0.01 |
|  | intra-center | 0.06 | 0.02 | 0.14 | 0.05 | 0.10 | 0.04 | 0.03 | 0.01 |
|  | inter-center | 0.02 | 0.01 | 0.10 | 0.04 | 0.07 | 0.03 | 0.01 | 0.01 |

However, when measured by SNVs in our defined truth set, cross-center and cross-platform variations were very small, indicating that all individual NGS runs, regardless of sequencing centers or NGS platforms, detected majority of “true” mutation consistently. This “consistency” dropped dramatically for SNVs not defined in the truth set, and further down to nearly zero, if SNVs haven’t been defined in the reference call set. For the purpose of equal coverage of each individual run in comparison group, read downsampling actually decreased the reproducibility of mutation detection. Both inter- and intra-center variations were much higher compared to results with original coverage.

Taken together, these results indicated that there were two major sources for non-concordant SNV calls between any two different library preparations: 1) stochastic effects of sequence coverage on SNVs with low MAF, which is sensitive to read coverage and mainly represented by SNVs in the group of “Not in-truth set”; 2) artifacts due to library preparation, mainly represented by SNVs in the group of “Not defined”. The benefit of high read coverage is not only to empower the detection of mutation with low MAF but also to increase result reproducibility (for both WES and WGS), probably due to reduced stochastic effect.

We repeated this analysis with results from MuTect2 and SomaticSniper (**Suppl. Table 5**). Results from the two callers not only confirmed the above observations but also supported conclusions from tornado plots and O_scores, i.e. for WES platform, Strelka2 on is less reproducible than MuTect2 (**Fig. 2c, Suppl. Fig. 4a, and Suppl. Fig. 6a**).

Using our established call set as a reference, we defined resulting SNV calls from any of NGS run with the pair of cell lines into three categories: 1) repeatable (SNVs in the truth set); 2) non-repeatable (SNVs not in the reference call set); and 3) gray-zone (SNVs not in the truth set) **(Suppl. Fig. 10)**. Regardless platforms or callers, the percentage of SNV in gray-zone was between 6-10%. While the repeatable calls in WGS was the highest (78%) for Strelka2, high number of Non-repeatable calls (61%) was also observed in WES runs. Repeatable calls by MuTect2 for both WES and WGS were quite high, 60% and 75%, respectively. In contrast, SomaticSniper had low number of repeatable calls (30-40%) and high number of non-repeatable calls (50-60%) for both WES and WGS platform, even though its repeatable calls in WES was higher than that in WGS. These results were consistent with what we have observed in tornado plot and O_Scores (**Suppl. Fig. 4a).**

The overall effect of each sequencing technique on the same replicate was also studied. When using Strelka2 as the caller, more unique mutations were called in WES than were identified in WGS (**Fig. 6a**). This remained true across different centers of replicate origin. Overall WGS and WES agreed, with correlation of MAFs from concordant SNVs ranging from 0.88-0.97 (**Fig. 6b**). Moreover, in cases with higher coverage, the concordance and correlation between WES and WGS were highest.

**Figure 6.**
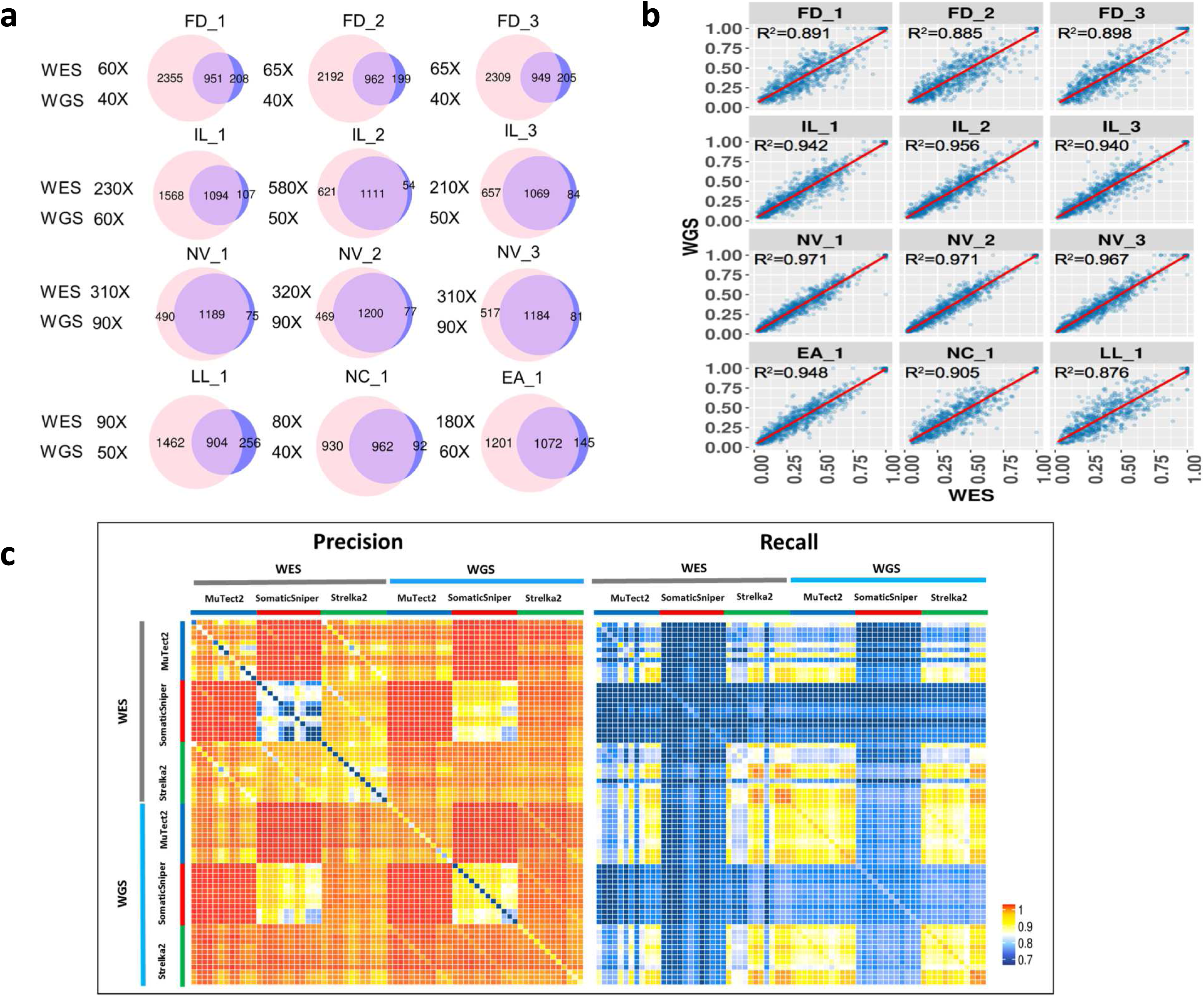
Comparison of concordance and correlation between results from WES and WGS with Strelka2 as the mutation caller. **(a)** SNVs/indels calling concordance between WES and WGS from twelve repeated runs. For direct comparison, SNVs/indels from WGS runs were limited to genomic regions defined by an exome capturing kit (SureSelct V6+UTR). WES is shown on the left in the Venn diagram and WGS is on the right. Shown coverage depths for WES and WGS were effective mean sequence coverage on exome region, i.e. coverage by total number of mapped reads after trimming. **(b)** Correlation of MAF in overlapping WGS and WES SNVs/indels from repeated runs. X- and Y-axis represents MAF detected in WES and WGS respectively. **(c)** Precision and recall of overlapping SNVs/indels that were supported by biological repeats (library repeats) or analytical repeats (two different callers). Each row or column represent calling results from a WES or WGS run called by one of the three callers from a BWA alignment.

The artifacts called by each sequencing type were then characterized. Variants were grouped into those common to WES and WGS, and those unique to one of the two platforms. The WES-only mutation calls were mostly characterized by lower MAF and higher C>A artifacts than were those in WGS (**Suppl. Fig. 11a, b**). Mutations identified by both WES and WGS had a more uniform distribution. Interestingly, the skew in artifacts in WES may be driven by library preparation protocol, where samples with shorter insert fragments were more skewed (**Fig. 1c**).

Precision and recall rates (**Fig. 6c**) from WES and WGS were also compared across all three callers and all twelve replicates directly. Mutations that were shared by two replicates generally had higher precision. Moreover, almost all mutations called by both MuTect2 and SomaticSniper were true. Leveraging additional callers increased precision; however, this was at the cost of recall (**Fig. 6c**). Taken together, by the precision metric, WGS clearly out-performed WES across replicates, callers, and sequencing centers. These figures demonstrate the importance of considering sufficient replicates during study design, rather than trying to compensate by using multiple callers.

Finally, we compared precision, recall, and F-score (as defined in Supplemental Methods) across a range of variant allele frequencies with three callers and in WES and WGS. Interestingly, Strelka2 exhibited the best performance on WGS samples but the worst performance on WES samples (**Suppl. Fig. 11c**), which was consistent with the results from the reproducibility study (**Fig. 2b-d**). On the other hand, even though SomaticSniper did not perform well overall, it was not affected by C>A artifacts. However, for both WES and WGS runs, we did observe that the lower limit of MAF calling by SomaticSniper was ∼12%, and many false positives were misidentified by SomaticSniper at a higher MAF (**Suppl. Fig. 12a, b**). Therefore, the “better” performance of SomaticSniper compared to Strelka2 on WES was driven by SomaticSniper’s insensitivity to artifacts from the DNA fragmentation process (**Fig. 1c and Suppl. Fig. 12a**).

## Discussion

In this study, we systematically evaluated the impact of several key steps on detecting cancer mutations with various NGS technologies. In addition to bioinformatics components such as preprocessing, alignment, post alignment, and mutation calling, we concomitantly queried the effects of biosamples, tumor purity, DNA input quantity, DNA fragmentation protocol, library preparation, and read coverage. Moreover, we investigated the reproducibility of WES and WGS across six laboratories, two NGS machines (HiSeq vs NovaSeq), three aligners, and three callers. Our results indicated that the reproducibility of cancer mutation detection not only depends on the choice of NGS platform (WGS vs. WES) but also on the selection of mutation callers (**Suppl. Fig. 4a**). Comparing to WGS, WES is very sensitive to sample processing, PCR effects, and choice of bioinformatics tools and thus less reproducible (**Fig. 2**).

Based on the results observed from this study, we note that each individual component of the sequencing and analysis process can have a tremendous impact on the final outcomes. For example, if FFPE samples were the original targets for inquiry, there would be artifacts from oxidative DNA damage. These artifacts pollute final mutation calls, but the level of artifact pollution is also dependent on NGS platform choice (WGS or WES), read preprocessing, and selected mutation caller. Similar artifacts also exist if DNA is over-fragmented by excessive sonication time. Less shearing time (**Fig. 1c**) or addition of antioxidant agents, such as ethylenediaminetetraacetic acid (EDTA), and butylated hydroxytoluene (BHT) in the sonication solution may reduce such artifacts^27^.

We also found that very low DNA input (1 ng) will require a PCR-based library preparation (such as TruSeq-Nano), resulting in a less diverse read population and consequently inferior detection capability. Of note, the relatively new Nextera Flex library preparation kit overcame the challenge of 1 ng DNA input by maintaining good library diversity, thereby achieving higher efficiency of mutation calling.

We concluded that read coverage is paramount to a robust analysis. Our results indicated that, to detect mutations with more than 5% MAF, read coverage between 50X-80X would be sufficient given samples with high tumor content (>50%); however, to detect low MAF mutations and/or mutations in samples with low tumor purity, high read coverage is essential. Researchers deciding the amount of coverage needed for given samples may reference our matrix table for actionable information (**Suppl. Table 4**). We recommend that users first assess the tumor content in each of their samples and then select read coverage accordingly. Should tumor content be low (<50%), enrichment of tumor purity, such as micro-dissection, would reduce the need for expensive high read coverage.

We took an alternative approach to evaluate the performance of various bioinformatics components and pipelines. The focus was on four key steps: preprocessing, alignment, post alignment, and mutation calling. By segregating these four sub-components, we were able to dissect the sources of discrepancies in SNV calling results obtained from multiple bioinformatics pipelines. Based on our results, we caution against using error-fixing procedures during read preprocessing steps to avoid introducing new artifacts while removing FFPE artifacts. Overall, we did not observe significant influence of alignments from three aligners: Bowtie2, BWA, and NovoAlign. However, the scale of alignment mapping quality from various aligners differed and consequently influenced mutation calling. To summarize, we saw a joint effect between aligner and caller, especially for callers optimized for a particular aligner. Thus, it is critical that the choice of caller and aligner should not be made in isolation since a strong interaction effect might be present.

Overall concordance and correlation of WES and WGS results were good. WES had a better coverage/cost ratio than WGS. However, the sequencing coverage on WES target regions was not even (**Suppl. Fig. 12c**). In addition, WES showed more batch effects/artifacts due to laboratory processing and thus had large variation between runs, laboratories and likely between researchers preparing the libraries. Overall run to run variations, including inter- and intra-center for WES were much higher than WGS (**Table 1**). As a result, WES was less reproducible than WGS. In comparison, WGS had more uniform coverage, was less sensitive to different runs and laboratories, and overall was more reproducible. Biological (library) repeats removed some artifacts due to random events and thus offered much better calling precision than did a single test. Analytical repeats (two bioinformatics pipelines) also increased calling precision at the cost of increased false negatives as well. Our results indicated that biological replicates are more important than bioinformatics replicates in cases where high specificity and sensitivity are needed.

Detection of cancer mutations is an inherently integrated process. No individual component can be singled out in isolation as more important than any others, and specific components interact with and affect each other. For example, a study aiming to detect mutation with >5% MAF in a cohort of FFPE 75% pure tumor biopsies should utilize 50X WGS coverage with MuTect2 calling on alignments from BWA. In cases where fresh frozen tissues are used with analysis on a WES platform, we recommend the use of the GIV (G>T/C>A) score to monitor the level of artifacts from DNA fragmentation processes and call mutations with Strelka2 on BWA alignments to detect mutations with <10% MAF and filter out false calls due to artifacts caused by DNA damage. Every component and every combination of components can lead to false discovery via both false positives and false negatives. Individual components of cancer mutation calling should never be considered “plug and play” given the effect one component has on all the other components in the mutation detection process. Although initial steps of NGS test validation may be performed in three separate stages (platform, test-specific, and informatics)^42^, our study demonstrates the complex interdependency of these stages on overall NGS test performance. Thus, final validation studies and revalidation when appropriate should be performed using the entire sample-to-result pipeline.

Until we finish similar studies on other cell lines, we cannot conclude that results from this study are generalizable to other cancer types. However, with high complexity of chromosome loss/gains (presented in another manuscript in preparation) and a large number of somatic mutations identified in HCC1395, this cell line resembled a hyperdiploid genome often seen in cancer. Our future direction is to expend the portfolio of reference samples/data sets by including other common cancer types, such lung cancer, colon cancer, and pancreatic cancer. Experience/knowledge accumulated via this study will provide guidelines for our future studies to be more efficient and cost-effective.

In summary, we investigated a series of NGS cancer mutation detection components, from wet lab to dry lab. The pair of tumor/normal cell lines, data sets with FFPE DNA, tumor/normal DNA mix, and performance of bioinformatics components established in this study can serve as a reference for the NGS research community to perform benchmarking studies for the development of new NGS products, assays, and informatics tools. To benefit the research community, we provided actionable recommendations on DNA fragmentation for WES runs, and a selection of NGS platforms, and bioinformatics tools based on the nature of available biosamples and study objectives.

## Methods

See Supplement for Methods

## Acknowledgements

We thank Sivakumar Gowrisankar of Novartis, Susan Chacko of the Center for Information Technology, the National Institute of Health for their assistance with data transfer, Dr. Jun Ye of Sentieon for providing the Sentieon software package. We also appreciate Dr. Laufey Amundadottir of the Division of Cancer Epidemiology and Genetics, National Cancer Institute (NCI), National Institutes of Health (NIH), for the sponsorship and the usage of the NIH Biowulf cluster; Dr Reena Phillip of the Center for Devices and Radiological Health, U.S. Food and Drug Administration, for her advices on study design; and Seven Bridges for providing storage and computational support on the Cancer Genomic Cloud (CGC). The CGC has been funded in whole or in part with Federal funds from the National Cancer Institute, National Institutes of Health, Contract No. HHSN261201400008C and ID/IQ Agreement No. 17×146 under Contract No. HHSN261201500003I. Research done in Dr. Charles Wang’s lab was partially supported by the NIH S10 grant, the Office of the Director, National Institute of Health under Award Number S10OD019960 (CW) and the Ardmore Institute of Health grant 2150141 (CW) and Dr. Charles A. Sims’ gift. This work also used the computational resources of the NIH Biowulf cluster (http://hpc.nih.gov). Original data was also backed up on the servers provided by Center for Biomedical Informatics and Information Technology (CBIIT), NCI.

In addition, we thank following people for their participation of working group discussions: Meredith Ashby, Ozan Aygun, Xiaopeng Bian, Pierre Bushel, Fabien Campagne, Tao Chen, Han-Yu Chuang, Youping Deng, Don Freed, Paul Giresi, Ping Gong, Yan Guo, Christos Hatzis, Susan Hester, Jonathan Keats, Eunice Lee, You Li, Sharon Liang, Tim McDianiel, Jai Pandey, Anand Pathak, Tieliu Shi, Jeff Trent, Mingyi Wang, Xiaojian Xu, Chaoyang Zhang.

## Disclaimer

This is a research study, not intended to guide clinical applications. The views presented in this article do not necessarily reflect current or future opinion or policy of the US Food and Drug Administration. Any mention of commercial products is for clarification and not intended as endorsement.

