## Supplementary Material for "Achieving reproducibility and accuracy in cancer mutation detection with whole-genome and whole-exome sequencing"

### Materials and methods

#### Cell line and DNA extraction

HCC1395; Breast Carcinoma; Human (*Homo sapiens*) cells (expanded from ATCC CRL-2324) were cultured in ATCC-formulated RPMI-1640 Medium, (ATCC 30-2001) supplemented with fetal bovine serum (ATCC 30-2020) to a final concentration of 10%. Cells were maintained at 37°C with 5% carbon dioxide ($CO_2$) and were subcultured every 2 to 3 days, per ATCC recommended procedures using 0.25% (w/v) Trypsin-0.53 mM EDTA solution (ATCC 30-2101), until appropriate densities were reached. HCC1395BL; B lymphoblast; Epstein-Barr virus (EBV) transformed; Human (*Homo sapiens*) cells (expanded from ATCC CRL-2325) were cultured in ATCC-formulated Iscove's Modified Dulbecco's Medium, (ATCC Catalog No. 30-2005) supplemented with fetal bovine serum (ATCC 30-2020) to a final concentration of 20%. Cells were maintained at 37°C with 5% $CO_2$ and were subcultured every 2 to 3 days, per ATCC recommended procedures, using centrifugation with subsequent resuspension in fresh medium until appropriate densities were reached. Final cell suspensions were spun down and re-suspended in PBS for nucleic acid extraction.

All cellular genomic material was extracted using a modified Phenol- Chloroform-Iso-Amyl alcohol extraction approach. Essentially, cell pellets were re-suspended in TE, subjected to lysis in a 2% TritonX-100/0.1% SDS/0.1 M NaCl/10mM Tris/1mM EDTA solution and were extracted with a mixture of glass beads and Phenol- Chloroform-Iso-Amyl alcohol. Following multiple rounds of extraction, the aqueous layer was further treated with Chloroform-IAA and finally underwent RNAse treatment and DNA precipitation using sodium acetate (3 M, pH 5.2) and ice cold Ethanol. The final DNA preparation was re-suspended in TE and stored at -80°C until use.

#### Cell line karyotyping

Karyotyping was performed by Cell Line Genetics (Madison, Wisconsin) essentially as described by Meisner LF et al[1]. Cells were treated with Colcemid (Gibco) for 40 min followed by exposure to 0.075M KCl for 23 min at 37$^{o}$C and then fixed with 3:1 methanol:glacial acetic acid. Slides were stained with Leishman's stain before observation. During observation, roughly 20 metaphase cells were counted at the microscope and numerical and structural chromosome aberrations were recorded. An analysis of 5–10 cells band for band was performed at the microscope using a 100x objective with an effort to karyotype at least two cells from each clone.

#### FFPE processing and DNA extraction

Cell lines cultured in T75 flasks (Corning Catolog No. 10-126-28) were harvested according to vendor product specifications (https://www.atcc.org/). For each cell line, the harvested materials were combined into a single 15 mL conical tube (Falcon Catalog No. 14-959-53A) and resuspended to a total volume of 1 mL with neutral buffered 10% formalin (StatLab Catalog No. 28600). In separate vials, HistoGel specimen processing gel matrix (ThermoFisher Catalog No. HG-4000-012) had been heated to 60°C for 2 hours to liquefy and then allowed to cool and equilibrate to 45°C in vendor supplied thermal block (ThermoFisher Catalog No. HGSK-2050-1). For each cell line, eight replicate rectangular shape cell-block molds were set up (Fisherbrand Catalog No. EDU00552). In each mold, 500 ul of 45°C HistoGel was added, and to this, 100 ul of neutral buffered formalin suspended cell line mixture was added. These were immediately and gently stirred to ensure homogeneity of cells within the cooling HistoGel matrix, and then allowed to sit and solidify on the bench top for at least 5 minutes. Next, for each mold, a micro-spatula was used to carefully dislodge the formed HistoGel embedded cell mixtures, and these were carefully placed into nylon mesh bags (Thermo Scientific Catalog No. 6774010) to prevent disaggregation during subsequent tissue processing. These formed HistoGel cell mixtures in nylon bags were placed into individual tissue processing cassettes (Thermo Scientific Catalog No. 1000957), and then submerged in plastic pail filled with neutral buffered 10% formalin to simulate pre-tissue processing time-in-formalin delay before batch tissue processing steps.

The sequence described above was performed at 1-, 2-, 6-, and 24-hour time points prior to batch tissue processing. All cassettes were then placed into a tissue processor for a “routine” tissue processing run at the University of Toledo Medical Center Department of Pathology (Sakura Tissue Tek VIP 5 Tissue Processor; see Supplementary Table for “routine” run conditions). The processed formalin fixed paraffin infiltrated cell blocks were then embedded in paraffin (Sakura Tissue Tek TEC 5 Tissue Embedding Station) to create formalin-fixed paraffin embedded (FFPE) cell blocks.

Each FFPE cell block was serially sectioned at 5 µm thickness with a microtome, and these ribbons of shaved material were placed into individual 15 ml conical tubes. A QIAamp DNA FFPE Tissue Kit (Qiagen, Duesseldorf, Germany) was used to extract FFPE DNA from each cell block following a slightly modified protocol. The first xylene step for deparaffinization was removed due to the low yield and purity of the DNA which is commonly experienced with clinical aspirate specimens or dyshesive specimens derived from cell culture specimens[2]. Instead, buffer ATL and proteinase K were directly added to the tubes with FFPE slices, and incubated according to vendor specifications. After digestion and lysis, the specimens were cooled to room temperature and continually inverted to let the paraffin solidify along the inside surface of the conical tubes. After cooling, the tubes were spun for 5 minutes at 1200 g and aqueous and paraffin layers become visible. The aqueous layer was carefully transferred to a new 15 ml conical tube. The specimens were then incubated at 90°C for 1 hour. The specimen was again cooled to room temperature and briefly centrifuged to remove liquid from the cap. The rest of the Qiagen QIAamp DNA FFPE Tissue Kit protocol, starting at the addition of Buffer AL and 100% ethanol with vortexing was followed according to vendor specifications. DNA was eluted from the QIAamp MinElute column using 100 ml of low concentration EDTA TE buffer (0.1 mM EDTA, Tris-HCl buffer, 10mM, pH8.5). Quality control for the specimens was performed using the following vendor instruments/kits: Thermo Scientific NanoDrop Spectrophotometry, Thermo Scientific Qubit fluorometer, absolute and relative qPCR measures of DNA quality, Agilent HighSensitivity D5000 Tapestation, and Agilent Highsensitivity DNA Bioanalyzer chip. A representative selection of the prepared cell blocks had a portion of their microtome sections taken for microscopic evaluation with hematoxylin and eosin (H&E) staining as well as immunohistochemistry. The routine H&E stained glass slides were used for estimates of cellularity, evenness of dispersion of cells in the cell block, and cytologic quality (viability and lack of degeneration in cellular membranes).

Immunohistochemistry for Pan Keratin (Ventana Catalog No. 760-2135) and CONFIRM-anti-CD45 (Ventana Catalog No. 760-2505) was performed using a Benchmark Ultra Ventana Automated IHC slide staining system. These two IHC stains were used to ensure that no cross-mixing of cellular materials occurred between the two cell lines during culture, harvesting and processing for FFPE (~10,000 cells assessed for each IHC staining/cell-line category).

#### DNA fragmentation and library preparation

The TruSeq DNA PCR-Free LT Kit (Illumina, FC-121-3001) was used to prepare samples for whole genome sequencing. DNA libraries for whole exome sequencing were firstly prepared with the Ovation Ultralow System V2 (NuGEN, 0347-A01) following the manufacturer's instructions. Then, exonic regions of each library (750 ng) were captured using the SureSelectXT Reagent kit (Agilent Technologies, G9611A), the SureSelectXT Human All Exon V6+UTR Capture Library (Agilent Technologies, 5190-8881) and the Ovation Target Capture Module (NuGEN, 0332-16) and following the manufacturer's instructions.

WGS libraries were prepared at six sites with the TruSeq DNA PCR-Free LT Kit (Illumina, FC- 121-3001) according to the manufacturers' protocol. One ug of DNA was used for the TruSeq-PCR-free libraries, unless specified otherwise. All sites used the same fragmentation conditions for WGS by using Covaris with targeted size of 350 bp. All replicated WGS and WES libraries were prepared on a different day. The input amount of WGS runs with fresh DNA was 1 ug unless otherwise specified. Detailed parameters of DNA fragmentation for twelve WES libraries are presented in **Suppl. Table 6**.

The concentration of the TruSeq DNA PCR-Free libraries for WGS was measured by qPCR with the KAPA Library Quantification Complete Kit (Universal) (Roche, KK4824). The concentration of all the other libraries was measured by fluorometry either on the Qubit 1.0 fluorometer or on the GloMax Luminometer with the Quant-iT dsDNA HS Assay kit (ThermoFisher Scientific, Q32854). The quality of all libraries was assessed by capillary electrophoresis either on the 2100 Bioanalyzer or TapeStation instrument (Agilent) in combination with the High Sensitivity DNA Kit (Agilent, 5067-4626) or the DNA 1000 Kit (Agilent, 5067-1504) or on the 4200 TapeStation instrument (Agilent) with the D1000 assay (Agilent, 5067-5582 and 5067-5583).

For the library preparation study, HCC1395 and HCC1395BL were diluted to 250, 100, 10 and 1ng in Resuspension Buffer (Illumina). For the 250 ng samples, libraries were generated using the Truseq DNA PCR-free protocol as described above. For the remaining samples, libraries were generated using the Truseq DNA Nano (Illumina) protocol as per manufacturer's instructions. DNA was sheared as described above, and the following PCR cycles were performed: 8 cycles for 100 ng input, 10 cycles for 10 ng input and 12 cycles for 1 ng input. Nextera Flex (Illumina) libraries were also prepared from 1, 10, and 100 ng inputs according to the manufacturer's instructions and amplified with 12, 8, and 5 cycles of PCR, respectively.

For the tumor purity study, 1µg tumor:normal dilutions were made in the following ratios using Resuspension Buffer (Illumina): 1:0, 3:1, 1:1, 1:4, 1:9, 1:19 and 0:1. Each ratio was diluted in triplicate. DNA was sheared using the Covaris S220 to target a 350 bp fragment size (Peak power 140w, Duty Factor 10%, 200 Cycles/Bursts, 55s, Temp 4 $^{o}$C). NGS library preparation was performed using the Truseq DNA PCR-free protocol (Illumina) following the manufacturer's recommendations.

For the FFPE study, SureSelect (Agilent) WES libraries were prepared according to the manufacturer's instructions for 200ng of DNA input, including reducing the shearing time to four minutes. Additionally, the adaptor-ligated libraries were split in half prior to amplification and one half amplified for 10 cycles and the other half for 11 cycles to ensure adequate yields for probe hybridization. Both halves were combined after PCR for the subsequent purification step. For WGS, NEBNext Ultra II (NEB) libraries were prepared according to the manufacturer's instructions. However, input adjustments were made according to the dCq obtained for each sample using the TruSeq FFPE DNA Library Prep QC Kit (Illumina) to account for differences in sample amplifiability. A total of 33 ng of amplifiable DNA was used as input for each sample.

#### DNA sequencing

Whole-genome libraries were sequenced on a HiSeq 4000 instrument (Illumina) at 2 x 150 bases read length with HiSeq 3000/4000 SBS chemistry (Illumina, FC-410-1003), and on a NovaSeq instrument (Illumina) at 2 x 150 bases read length using the S2 configuration (Illumina, PN 20012860). Whole-exome libraries were sequenced on a HiSeq 2500 instrument (Illumina) at 2 x 125 bases read length and using HiSeq Rapid SBS v2 chemistry (Illumina, FC-402-4021 and FC-402-4022). In all cases, sequencing was performed following the manufacturer's instructions.

FASTQ sequence files for whole-genome and whole-exome sequencing were generated from the Illumina sequencer images using the Illumina RTA 1.18.66.3 (HiSeq 2500) or 2.7.7 (HiSeq 4000) and bcl2fastq 2.17.1.14 software.

#### Read processing and quality assessment

The FASTQ files generated for whole-genome and whole-exome sequencing from each sequencing center were transferred to central storage for read preprocessing and quality control. The Illumina bcl2fastq2 (v2.17) was used to demultiplex and convert binary base calls and qualities to FASTQ format. FASTQC(v0.11.2 )[3] was run on the raw reads to assess basecall quality, adapter content, G/C content, sequencing length and duplication level. In addition, FASTQ_screen(v0.5.1) and miniKraken(v0.10.0)[4] were run to detect possible cross contamination with other species. A multiQC(v1.3) run report was generated for each sample set. The sequencing reads were trimmed of adapters and low quality bases using Trimmomatic(v0.30)[5]. The trimmed reads were mapped to the human reference genome GRCm38 (see the read alignment section) using BWA-mem(v0.7.12)[6] in paired-end mode. In addition, the DNA Damage Estimator(v3)[7] was used to calculate the GIV score based on an imbalance between R1 and R2 variant frequency of the sequencing reads to estimate the level of DNA damage that was introduced in the sample/library preparation processes. Post alignment QC was performed based on BWA alignment BAM files, the genome mapped percentages and mapped reads duplication rates calculated by BamTools (v2.2.3) and Picard (v1.84)[8]. The genome coverage and exome target region coverages as well as mapped reads insert sizes, and G/C contents were profiled using Qualimap(v2.2)[9] and custom scripts. Preprocessing QC reports were generated during each step of the process. MultiQC(v1.3)[10] was run to generate an aggregated report in html format. A standard QC metrics report was generated from a custom script.

In order to assess trimming and error correction effects on the mutation call precision and recall, we chose a trimming software tool (Trimmomatic) and an error correction software package (BFC). For Trimmomatic, we used "MAXINFO: 50:0.97" to run against the same set as WES for benchmarking. BFC version 1.0-7-g69ab176[11] was run with default parameters aside from k value to provide corrected reads. Since BFC has an upper limit of 62 for k, FASTQ files with a larger optimal k value were processed with k=62.

#### Read alignment

For all alignments, we used the decoy version of the hg38 human reference genome (https://gdc.cancer.gov/about-data/data-harmonization-and-generation/gdc-reference-files; GRCh38.d1.dv1.fa) utilized by the Genomic Data Commons (GDC). For alignment comparisons, we ran NovoAlign v3.07.01 (Novocraft Technologies), Bowtie2 v2.2.9[12], and BWA-MEM v0.7.17[13]. Bowtie2 was run using all default parameters and bwa-mem was run with the –M flag for downstream Picard compatibility. Due to the prohibitively slow speed of NovoAlign, and to improve multithreading performance, we split each sample's reads into 20 separate batches of equal size, and then mapped each batch of 20 using 32 threads with NovoAlign and default parameters.

#### Read downsampling and pooling

Sequencing reads were downsampled using SAMtools version 1.6 on the BioGenLink™ platform ("BGL"; https://biogenlink.atlassian.net/wiki/spaces/BD/overview). A workflow was created in BGL called "Multi downsample BAM", which runs the "SAMtools view" tool on all SAM or BAM files in a directory and includes an option to downsample the reads by a given fraction corresponding to the "-s" parameter in SAMtools view. The workflow indexed the resulting BAM files using "SAMtools index". The workflow was used to generate all downsampled BAM files and index files and created a subset with defined read coverage.

BAM files from BWA[6] alignment of three replicated runs of WGS with 100X coverage on HCC1395 and HCC1395BL were merged using SAMtools (version 1.8)[14] for 200X or 300X coverages respectively. Newly created BAM files were then indexed and regrouped using Picard Tools (version 2.17.11)[8].

#### Assessment of reproducibility and O_Score calculation

We created and used "tornado" plots to visualize the consistency of mutation calls derived from aligners, callers, or repeated NGS runs. The height of the "tornado" represents the number of overlapping calls in the VCF files in descending order. The top of each plot portrays SNVs called in every VCF file. The bottom of the plots contains SNVs present in only one VCF file. The width of the "tornado" represents the number of accumulated SNVs in that overlapping category, which is scaled by the total number of SNVs in the corresponding sub-group. In addition, we established following formula to measure reproducibility based on the overlapping SNVs:

$$O_{score} = \frac{\sum_{i=1}^{i \to n}((\frac{i}{n}) \times O_i)}{\sum_{i=1}^{i \to n} O_i}$$

where *n* is the total number of VCF results in the pool set, $i$ is the number of overlaps, $O_i$ is the number of accumulated SNVs in the set with $i$ number of overlapping**.**

Statistical analysis was performed to evaluate the sources of variances in WES and WGS O_Scores (JMP Genomics 9.0). For WES, a primary fixed effect linear regression was first used to screen for 2-degree interaction terms significantly contributing to the outcome (F-test P<0.05). All possible 2-degree interactions, along with original variables were included in this primary model, and 5 interaction terms (Callers*Mean Coverage Depth, Callers*Percent GC, Machine model*Callers, Callers*GIV(G>T), and Callers*Percent Non-duplicated Reads) were significant (P<0.05). A linear transform was applied to individual variables to rescale the data to range from -1 to 1. The final fixed effect linear regression for WES included a total of 13 variables (8 original variables and 5 interactions). For WGS, we did not include any interactions as the individual variables accounted for >99% of the O_Score variance. The coefficient of determination ($R^2$) was calculated for both models. In addition, we calculated F statistics and corresponding P values for the variables included in the final model to measure their effects on the O_Score. Pairwise Pearson correlation coefficients between continuous variables were also calculated for both platforms.

#### Analysis of inter-/intra-center variations for WES and WGS

We performed the following analysis to further investigate inter- and intra-center reproducibility based on concordance of SNV detection in any pair of NGS runs, in association of SNVs defined in our call set, as described in section for “Creating a high-confidence call truth set”. We used the Jaccard index[15] to measure the concordance of SNVs from any “non-self” pair of runs:

$$J(A,B) = \frac{|A \cap B|}{|A \cup B|} = \frac{|A \cap B|}{|A| + |B| - |A \cap B|}.$$

A total of four comparison groups were included in this analysis: a) WES (12 repeats/3 centers), b) WGS (limited to exome regions, 12 repeats/3 centers), c) WGS (12 repeats/3 centers), and d) WES vs WGS (limited to exome regions, 24 repeats/3 centers). We did each of the four comparison groups with “all reads” (original coverage) and “downsampled reads” (fixed coverage of each NGS run, 50X for WGS and 150X for WES).

In addition, to investigate reproducibility represented by overall SNVs called in a given pair of NGS runs, we also broke down SNVs into three subgroups: a) In-truth set: SNVs defined in “HighConf” and

“MedConf” categories in the call set; b) Not in-truth set: SNVs defined in “LowConf” and “Unclassified” categories in the call set; and c) Not defined: SNVs not defined in the call set. Jaccard scores for any pair of NGS runs were calculated as depicted in the following figure and formulas based on: 1) overall SNV; 2) In-truth set; 3) Not in-truth set; and 4) Not defined.

Average value of Jaccard scores from all possible pairs of NGS runs in each of four comparison groups was aggregated at groups for “Overall pairs”, “intra-center”, or “inter-center”.

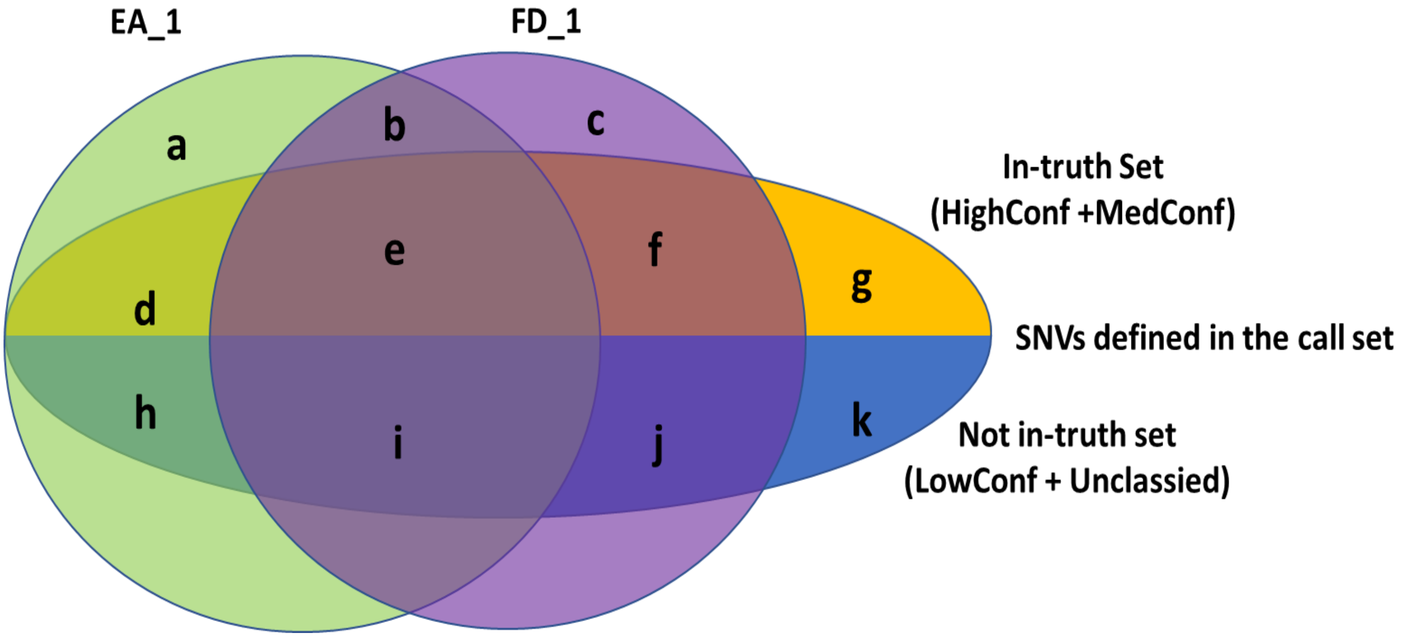


$$J(\text{Overall SNV}) = \frac{(a+b+d+e+g+i) \cap (b+c+e+f+g+h)}{(a+b+d+e+g+i) \cup (b+c+e+f+g+h)} = \frac{(b+e+i)}{(a+b+c+d+e+f+g+h+i)}$$

$$J(\text{In-truth set}) = \frac{(d+e) \cap (e+f)}{(d+e) \cup (e+f)} = \frac{e}{(d+e+f)}$$

$$J(\text{Not in-truth set}) = \frac{(h+i) \cap (i+j)}{(h+i) \cup (i+j)} = \frac{i}{(h+i+j)}$$

$$J(\text{Not defined}) = \frac{(a+b) \cap (b+c)}{(a+b) \cup (b+c)} = \frac{b}{(a+b+c)}$$

#### Somatic SNV callers

We used four somatic variant callers, MuTect2 (GATK 3.8-0)[16], SomaticSniper (1.0.5.0)[17], Lancet (1.0.7), and Strelka2 (2.8.4)[18] that are readily available on the NIH Biowulf cluster, and ran each of them using the default parameters or parameters recommended by the user's manual. Specifically, for MuTect2, we included flags for "-nct 1 -rf DuplicateRead -rf FailsVendorQualityCheck -rf NotPrimaryAlignment -rf BadMate -rf MappingQualityUnavailable -rf UnmappedRead -rf BadCigar", in order to avoid the running exception for "Somehow the requested coordinate is not covered by the read". For MuTect2, we used COSMIC v82 as required inputs. For SomaticSniper, we added a flag for "-Q 40 -G -L –F", as suggested by its original author, to ensure quality scores and reduce likely false positives. For TNscope (201711.03), we used the version implemented in Seven Bridges's CGC with the following command, "sentieon driver -i $tumor_bam -i $normal_bam -r $ref --algo TNscope --tumor_sample $tumor_sample_name --normal_sample $normal_sample_name -d $dbsnp $output_vcf". For Lancet, we ran with 24 threads on the following parameters "--num-threads 24 --cov-thr 10 --cov-ratio 0.005 --max-indel-len 50 -e 0.005". Strelka2 was run with 24 threads with the default configuration. The rest of the software analyzed was run as a single thread on each computer node. All mutation calling on WES data was performed with the specified genome region in a BED file for exome-capture target sequences. The high confidence outputs or SNVs flagged as "PASS" in the resulting VCF files were applied to our comparison analysis. Results from each caller used for comparison were all mutation candidates that users would otherwise consider as "real" mutations detected by this caller.

The performance of SNV callers was compared using the following metrics:

recall = # of true positives/(# of true positives + # of false negatives);

precision = # of true positives/(# of true positives + # of false positives);

F-score = 2 x (precision x recall)/(precision + recall);

Additional information is provided in the supplementary materials.

#### GATK indel realignment and quality score recalibration

The GATK (3.8-0)-IndelRealigner was used to perform indel adjustment with reference indels defined in the 1000Genome project (https://www.google.com/url?sa=t&rct=j&q=&esrc=s&source=web&cd=4&ved=0ahUKEwjlkcfB5-nbAhVOhq0KHXUWCKUQFgg7MAM&url=ftp%3A%2F%2Fftp.1000genomes.ebi.ac.uk%2Fvol1%2Fftp%2Ftechnical%2Freference%2FGRCh38_reference_genome%2Fother_mapping_resources%2FALL.wgs.1000G_phase3.GRCh38.ncbi_remapper.20150424.shapeit2_indels.vcf.gz&usg=AOvVaw0pLCj6zDgJg0A6zbFeMfQl). The resulting BAM files were then recalibrated with regard to quality with "BaseRecalibrator" and dbSNP build 146 as the SNP reference. Finally, "PrintReads" was used to generate recalibrated BAM files.

#### Creating a high-confidence call truth set

The high-confidence somatic SNV/INDEL truth set was created based on the reproducibility of the five Data Groups that included 63 pairs of WGS BAM files following these basic steps.

1. Use GATK CombineVariants to merge 63 SNV VCF files with 63 INDEL VCF files (54 of which were classified by SomaticSeq classifiers and 9 classified by majority-vote consensus).
2. Use SomaticSeq:seqc2_v0.4 to assign initial tiers based on cross-institution and cross-aligner reproducibility of each variant.
    a. For each call, use SomaticSeq:seqc2_v0.4 to extract VAFs from the 300X tumor-normal titration data sets and assess whether the VAF moves as expected with titration.
    b. For each call, use SomaticSeq:seqc2_v0.4 to extract sequencing features from each of the 63 x 2 = 126 BAM files.
3. Based on the initial tiers, sequencing features, and additional validation data, annotate the confidence level for each variant to create the somatic mutation truth set.
4. Exclude SNVs/indels in chrX, the short-arm of chr6 and the long arm of chr16 from the truth set since cytogenetic analysis has indicated their loss in HCC1395BL (**Suppl. Fig. 8b**).

The SomaticSeq pipeline was implemented and run on Seven Bridges' Cancer Genomics Cloud (CGC).

#### Data availability

All raw data (FASTQ files) are available on NCBI's SRA database (SRP162370). The truth set for somatic mutations in HCC1395, VCF files derived from individual WES and WGS runs, and source codes for "tornado" plot are available on NCBI's ftp site (ftp://ftp-trace.ncbi.nlm.nih.gov/seqc/ftp/release/Somatic_Mutation_WG/). Some alignment files (BAM) are also available on Seven Bridges' s Cancer Genomics Cloud (CGC) platform and Digicon's BioGenLink™.

### Supplementary Figures

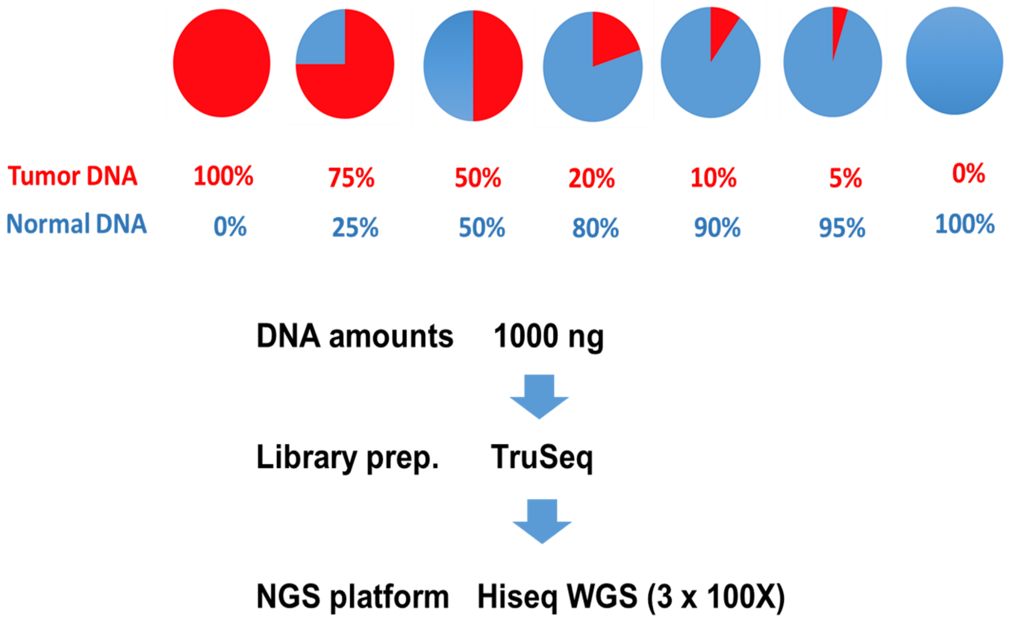


**Supp. Fig.1**. Tumor purity study experiment design. 1000 ng of HCC1395:HCC1395BL dilutions were made in the following ratios: 1:0, 3:1, 1:1, 1:4, 1:9, 1:19 and 0:1. Each ratio was diluted in triplicate. NSG library preparation was performed using the TruSeq DNA PCR-free manufacturer's protocol (Illumina) following the manufacturer's recommendations. Libraries were sequenced to ~3 billion reads by pooling two samples per flowcell using the paired end 150 base single index read format as per manufacturer's recommendations.

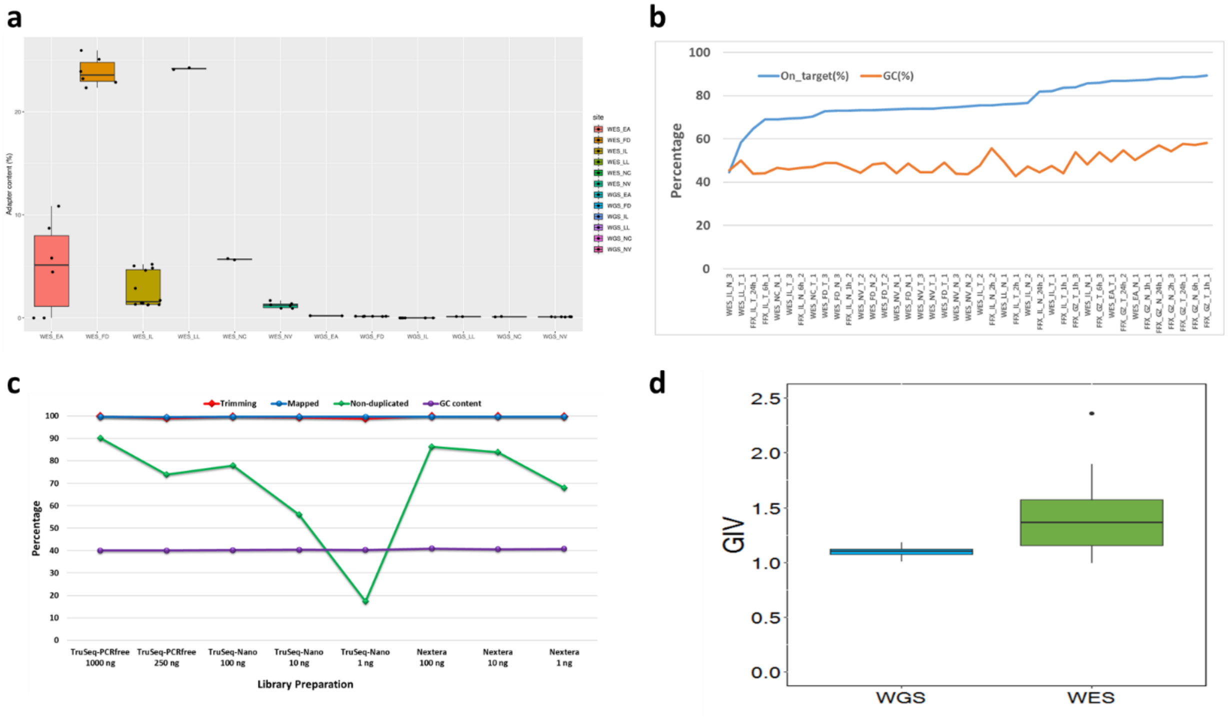


**Supp. Fig.2**. Read mapping quality statistics. **(a)** Adapter sequence content in WES and WGS libraries. **(b)** Percentage of reads mapped to target regions (SureSelect V6+UTR) and GC content for WES runs on fresh or FFPE DNA. **(c)** Read quality from three WGS library preparation kits (TruSeq PCRfree, TruSeq-Nano, and Nextera Flex) on fresh or FFPE DNA. **(d)** Distribution of GIV scores in WGS and WES runs.

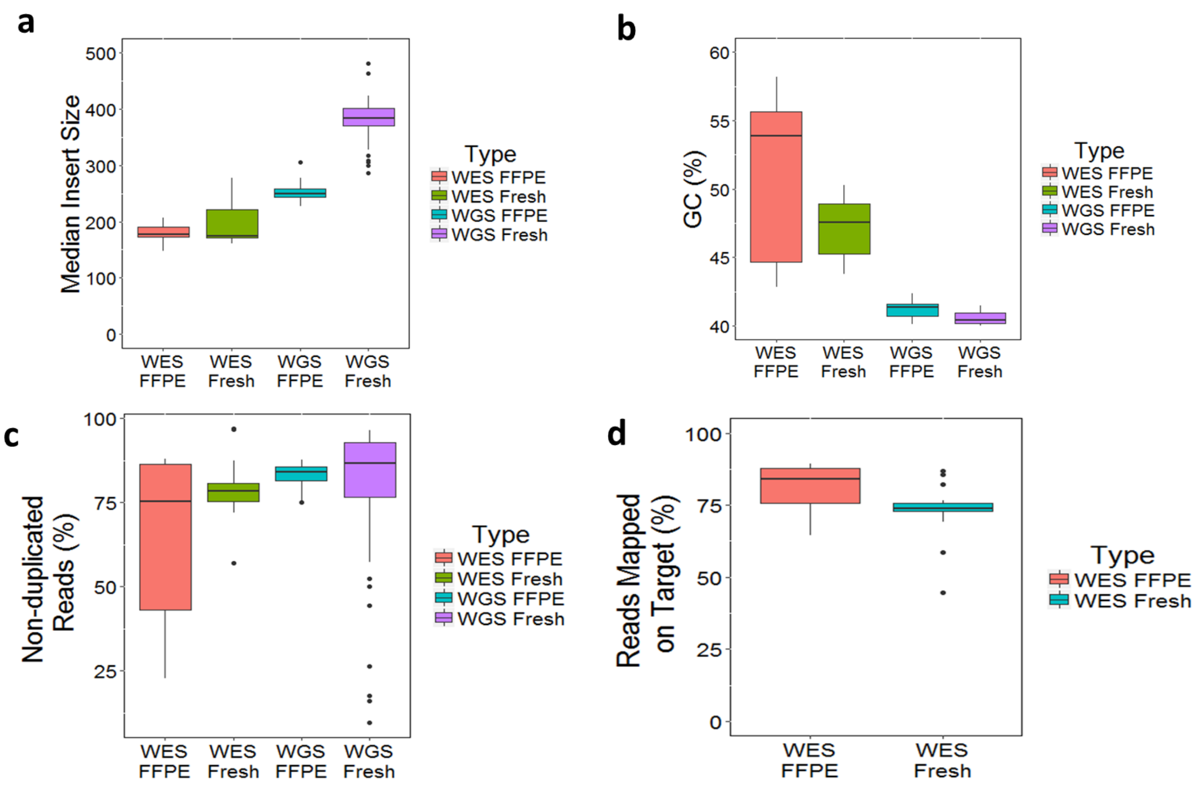


**Supp. Fig.3**. Overall read quality distribution for all WES and WGS runs. **(a)** Median insert fragment size of WEG and WGS run on fresh and FFPE DNA. **(b)** G/C content of reads for WES and WGS runs. **(c)** Overall read redundancy for WES and WGS runs. Some outliers were observed in WGS fresh DNA, which were from runs of TruSeq-Nano with 1 ng of DNA input. **(d)** Overall percentage of reads mapped to target regions for WES runs for fresh and FFPE DNA.

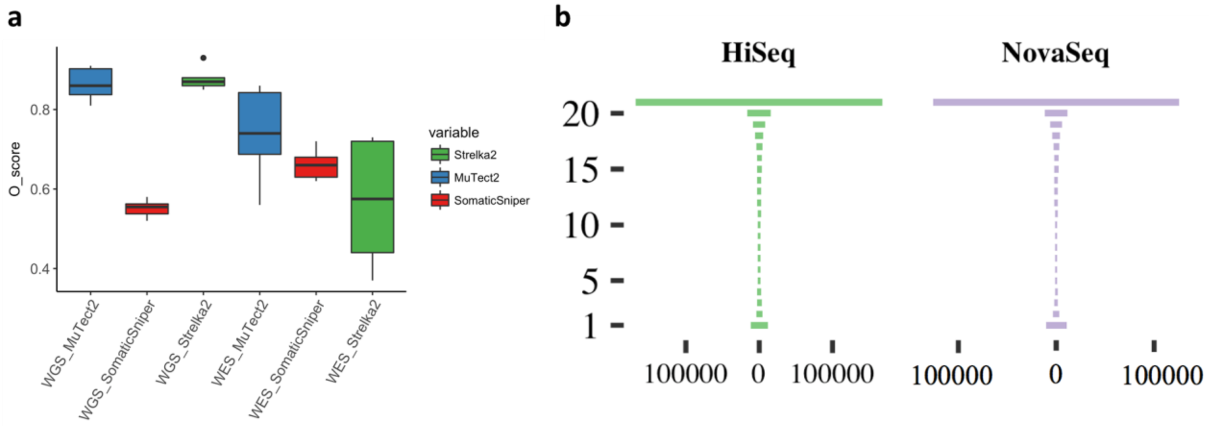


**Supp. Fig. 4**. Mutation calling repeatability and O_Score distribution. **(a)** Distribution of O_Score by three callers (MuTect2, Strelka2, and SomaticSniper) for 12 WGS and WES runs on BWA alignments. **(b)** "Tornado" plot of reproducibility between 12 WGS runs on HiSeq series (2500, 4000, X10) and 9 WGS runs on NovaSeq (S6000). SNVs/indels were called by Strelka2 on BWA alignments.

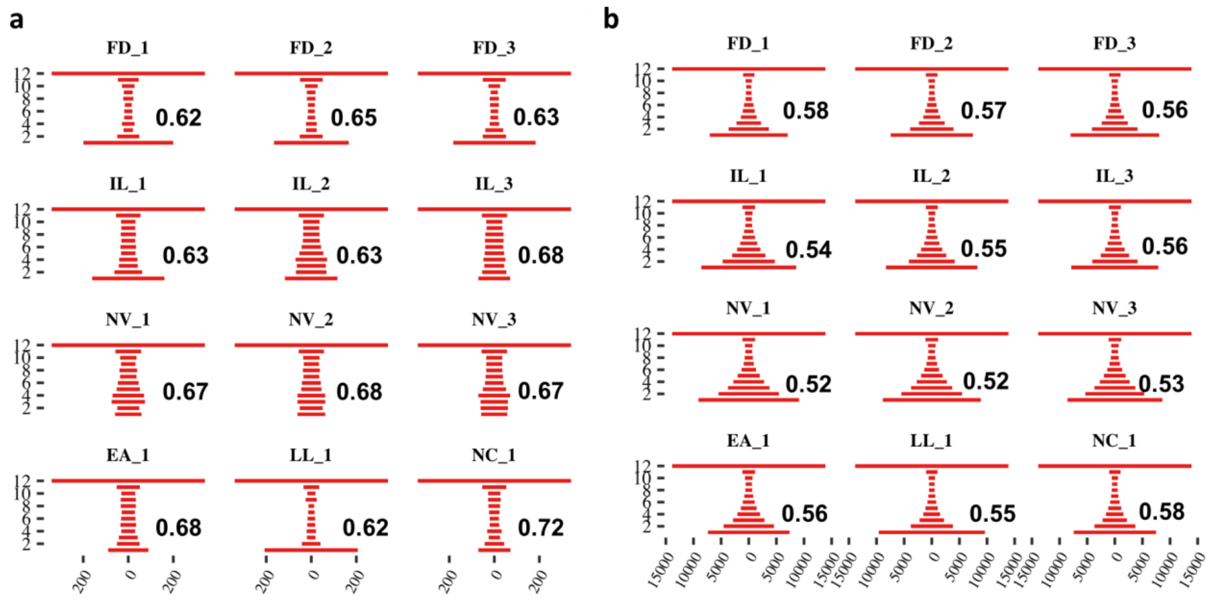


**Supp. Fig. 5**. Mutation calling repeatability by SomaticSniper on BWA alignments. "Tornado" plots and O_Scores for 12 WES repeats **(a)** and 12 WGS repeats **(b)**.

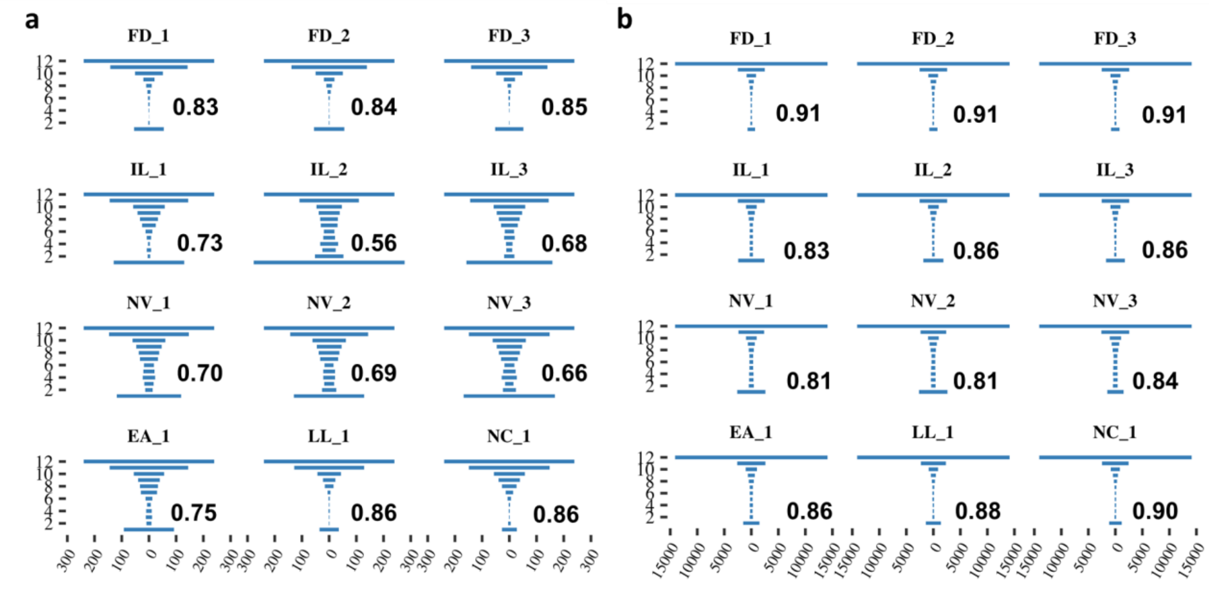


**Supp. Fig. 6**. Mutation calling repeatability by MuTect2 on BWA alignments. "Tornado" plots and O_Scores for 12 WES repeats **(a)** or 12 WGS repeats **(b)**.

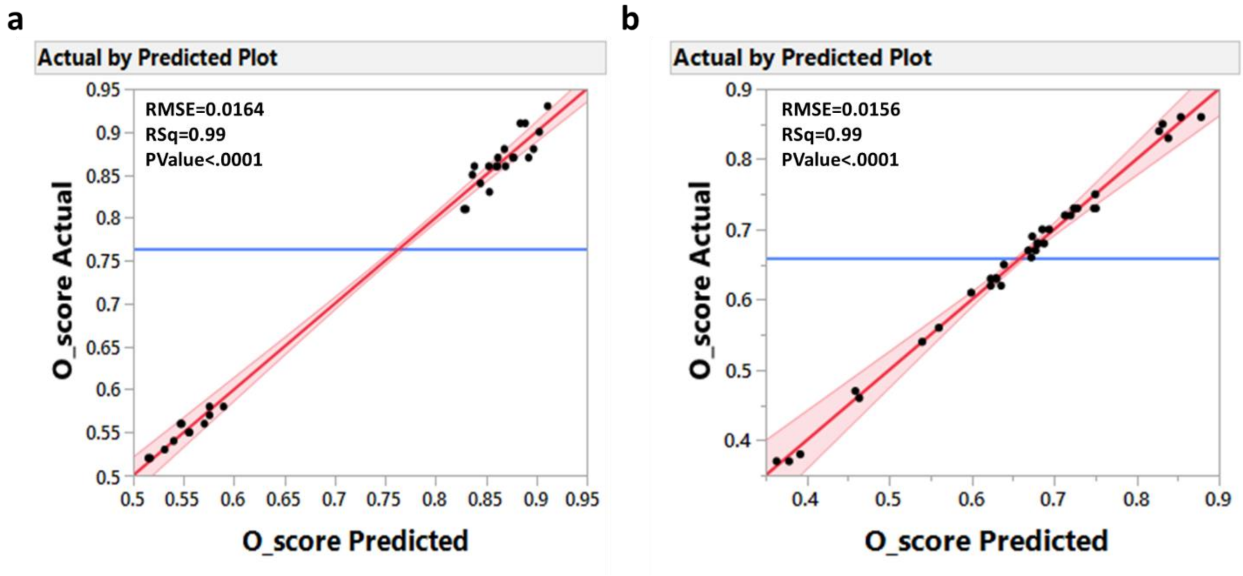


**Supp. Fig. 7**. Source of variance in reproducibility measured by O_Score. Actual by Predicted plot of WGS (a) and WES (b). A total of 8 variables (WGS) or 13 variables (WES), including 2-degree interactions, were included in the fixed effect linear model.

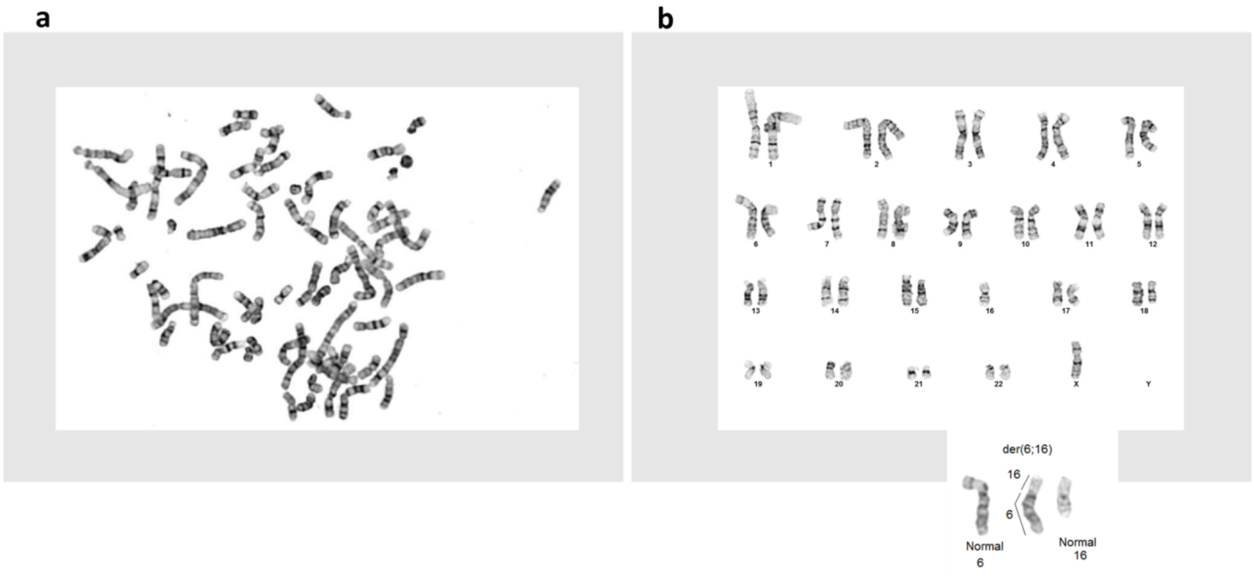


**Supp. Fig. 8**. Karyotyping of HCC1395 and HCC1395BL. **(a)** Karyotype of HCC1395. Cytogenetic analysis was performed on ten G-Banded metaphase cells from HCC1395. It showed as a hypertetraploid line with chromosome counts ranging from 64-79 and gain of 38-63 unidentifiable marker chromosomes. **(b)** Karyotype of HCC1395BL. Cytogenetic analysis was performed on ten G-banded metaphase cells from HCC1395BL. All ten cells showed loss of a chrX and an unbalanced whole arm translocation between the long-arm of chr6 at band q10 and the short-arm of chr16 at band p10. This results in a net loss of one copy of the short-arm of chr6 and loss of one copy of the long-arm of chr16. The abnormal chromosome could be placed in either a chr6 or chr16 loci as we were unable to determine if the centromere belongs to chr6 or chr16 (inset figure).

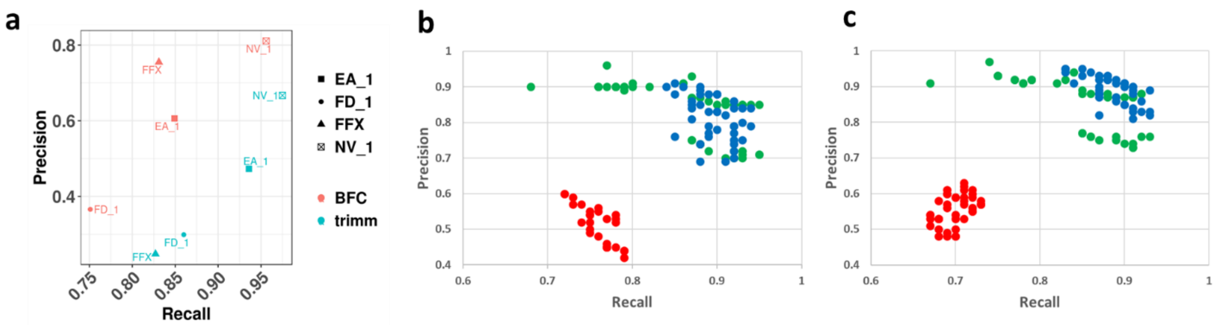


**Suppl. Fig. 9**. Impact of post alignment processing on precision and recall of WES and WGS run on FFPE DNA. **(a)** Precision and recall of mutation calls by Strelka2 on BWA alignments. A single library of FFPE DNA (FFX) and three libraries of fresh DNA (EA_1, FD_1, and NV_1) on a WES platform. Resulting reads were either processed by BFC tool or by Trimmomatic. Processed fastq files were then aligned by BWA and called by strelka2. Precision and recall were derived by matching calling results with truth set. Precision and recall of mutation calls by three callers, MuTect2 (blue color), Strelka2 (green color), and SomaticSniper (red color), on BWA alignments without **(b)** or with **(c)** GATK post alignment process (indel realignment & BQSR).

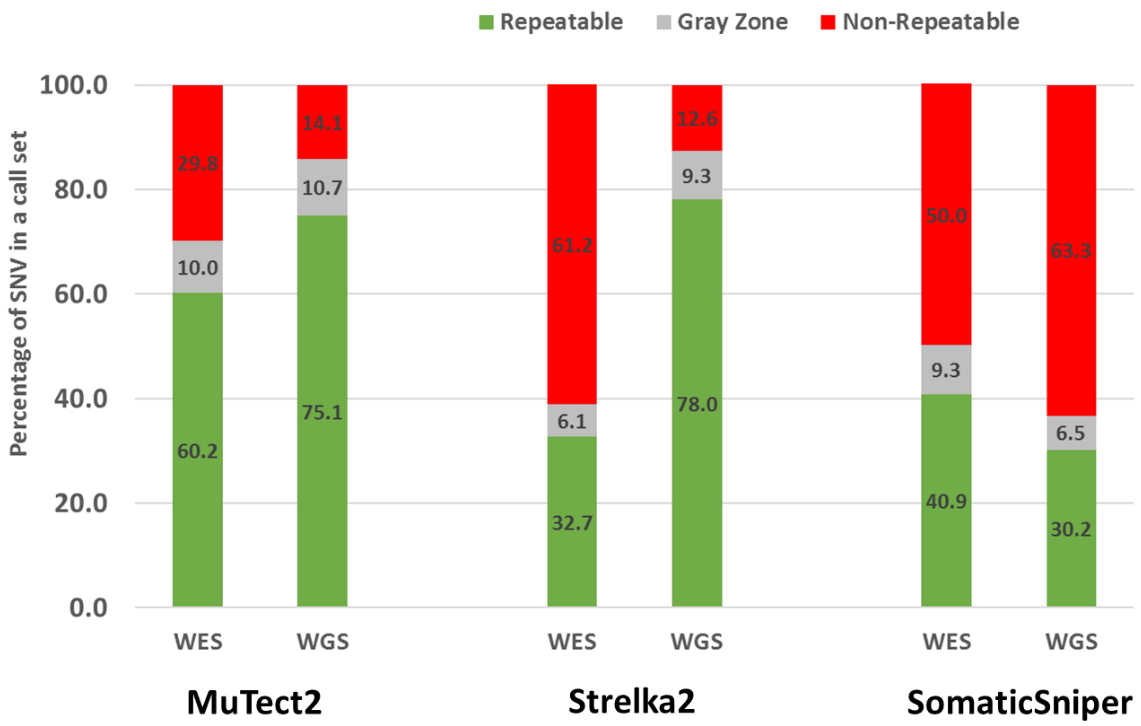


**Supp. Fig. 10**. Percentage of SNVs in a call set of WGS or WES run by three callers. Repeatable: SNVs defined in the truth set of the reference call set; Gray-zone: SNVs not defined as “truth” in the reference call set; Non-repeatable: SNVs were not in the reference call set.

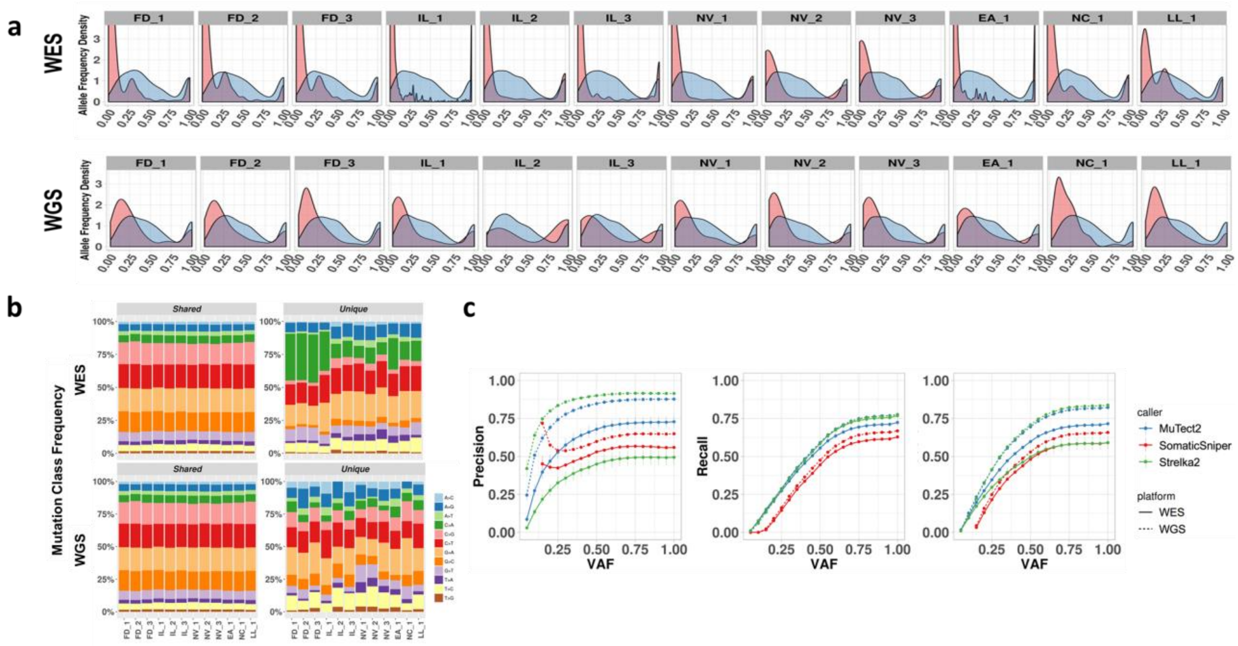


**Supp. Fig. 11**. WGS vs. WES platform-specific mutations and allele frequency calling accuracy. **(a)** Distribution of MAF for platform-only (shaded in pink) or shared SNV (shaded in blue) mutations by 12 repeated WES and WGS runs. X-axis: Mutation allele frequency (MAF) **(b)** percentage of mutation types for both WES (upper panel) and WGS (lower panel), in either shared (left) or unique (right). Note, high percentage of C/A mutation in the sites that also have high G/T_C/A GIV scores as displayed in **Fig. 1d**. **(c)** Cumulative MAF plot of precision, recall, and F-Score for three callers (MuTect2, Strelka2, and SomaticSniper) on WES and WGS runs.

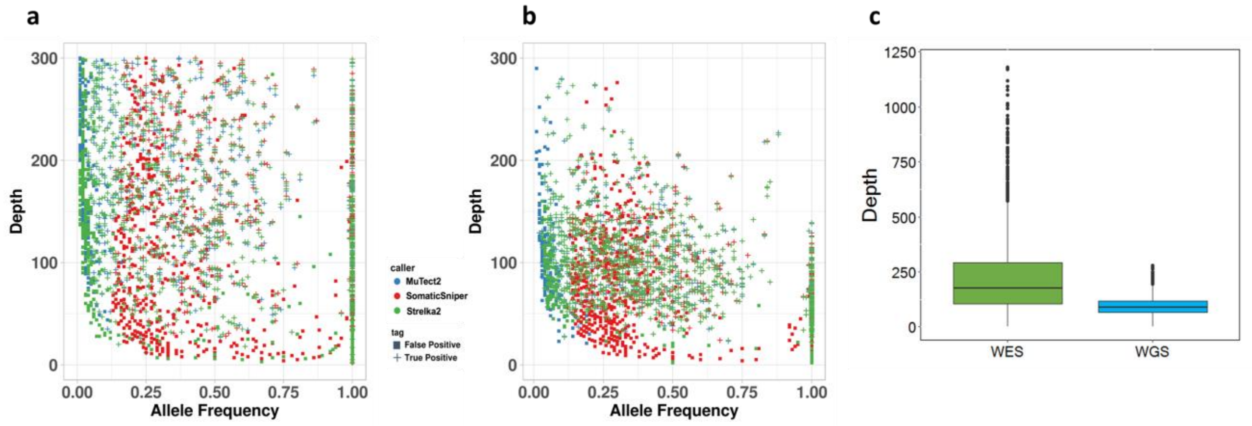


**Supp. Fig. 12**. Mutation allele frequency and coverage depth in WES and WGS sample. Scatter plot of allele frequency and their coverage depth by three callers, MuTect2, Strelka2, and SomaticSniper in one example WES sample **(a)** or WGS sample **(b). (c)** Boxplot of reads depth on called mutations in WES or WGS.
